# Inositol polyphosphate multikinase regulates Th1 and Th17 cell differentiation by controlling Akt-mTOR signaling

**DOI:** 10.1101/2024.01.08.574595

**Authors:** Chae Min Yuk, Dongeon Kim, Sehoon Hong, Mingyo Kim, Hyun-Woo Jeong, Seung Ju Park, Hyungyu Min, Wooseob Kim, Sang-Gyu Kim, Rho Hyun Seong, Seyun Kim, Seung-Hyo Lee

## Abstract

Activated proinflammatory T helper (Th) cells, such as Th1 and Th17 cells, mediate immune responses against intra- and extra-cellular pathogens as well as cause the development of various autoimmune diseases. Inositol polyphosphate multikinase (IPMK) is a key enzyme essential for inositol phosphate and phosphoinositide metabolism, which is known to control major biological events such as growth; however, its role in the function of Th cells remains unclear. Here we show that the expression of IPMK is highly induced in distinct Th1 and Th17 subsets. Further, while conditional deletion of IPMK in CD4^+^ T cells is dispensable for Th2-dependent immune responses, both Th1- and Th17-mediated immune responses are markedly diminished when this enzyme is absent resulting in reduced resistance to *Leishmania major* infection and attenuation of experimental autoimmune encephalomyelitis (EAE), an animal model of multiple sclerosis. In addition, IPMK-deficient naive CD4^+^ T cells display aberrant T cell activation and impaired differentiation into Th17 cells, which is associated with reduced activation of Akt, mechanistic target of rapamycin (mTOR), and STAT3. Mechanistically, IPMK as a phosphatidylinositol 3-kinase (PI3-kinase) controls the production of phosphatidylinositol (3,4,5)-trisphosphate, thereby promoting T cell activation, differentiation, and effector functions. Our findings suggest that IPMK acts as a critical regulator of Th1 and Th17 differentiation, highlighting the physiological importance of IPMK in Th1- and Th17-mediated immune homeostasis.

## Introduction

CD4^+^ T helper (Th) cells not only play critical roles in mediating adaptive immune responses to various pathogens, but are also involved in allergic responses, autoimmunity, and tumor immunity. Naive CD4^+^ T cells become effector and/or memory cells with specialized phenotypes, including Th1, Th2, Th17, and regulatory T (Treg) cells, as defined by signature cytokines and master transcription factors. Interleukin (IL)-12 and interferon (IFN)-γ are major inducers of Th1 cells, while the master transcriptional regulator for Th1 cells is T-bet. While induction of T-bet is dependent on signaling transducer and activator of transcription 1 (STAT1), IL-12 signaling activates STAT4. IL-4 is a critical cytokine for Th2 cell differentiation, and IL-4-induced STAT6 upregulates the expression of the Th2 cell master regulator, GATA3. Th17 cells develop in the presence of IL-6, TGF-β, IL-21, and IL-23, which induce the upregulation of the master regulator retinoic acid receptor-related orphan gamma T (RORγt) through STAT3. In addition, Treg cells express the master transcription factor, Foxp3, and develop in peripheral lymphoid organs after antigen priming (iTreg) or in the thymus (nTreg) (Luckheeram, Zhou et al. 2012).

Th1 cells are required for the immune responses against intracellular pathogens, such as viruses and intracellular bacteria, and exert their effects through the secretion of IFN-γ, which activates macrophages for enhanced phagocytic function. However, uncontrolled Th1-mediated responses cause autoimmunity by inducing chronic inflammatory responses as well as Th17 cells. Th17 cells, characterized by the production of signature cytokines IL-17A, IL-17F and IL-22, are thought to be critically involved in infection, inflammation, autoimmunity, and cancer (Dong 2008, Zou and Restifo 2010, Peters, Lee and Kuchroo 2011). Dysregulated Th17 cells play roles in the development of various autoimmune diseases, including psoriasis, inflammatory bowel disease, rheumatoid arthritis, and multiple sclerosis (Korn, Bettelli et al. 2009). Further, STAT3, activated by IL-6, functions as a promoter of *Rorc, Il17a, Il17f* and multiple other genes associated with Th17 cell polarization and survival (Durant, Watford et al. 2010). Consequently, IL-17 causes granulopoiesis by inducing granulocyte-colony-stimulating factor (GM-CSF) secretion in the stroma of bone marrow and promotes the expression of TNF-α, IL-1β, IL-6, chemokines and the GM-CSF receptor (Fossiez, Djossou et al. 1996, Tan, Huang et al. 2006). Therefore, IL-17 is highly pleiotropic and induces a range of inflammatory effects that link innate and adaptive immunity. As a result, Th17 cells cause tissue damage by recruiting neutrophils and macrophages into peripheral tissues.

Upon antigen presentation, T cell receptors (TCRs) and costimulatory molecules trigger signaling pathways including the activation of phosphatidylinositol 3-kinase (PI3K), which leads to the phosphorylation on the third position of phosphatidylinositol(4,5)-bisphosphate (PIP_2_) to produce phosphatidylinositol (3,4,5)-trisphosphate (PIP_3_), a potent second messenger known to promote Akt activities (Kane and Weiss 2003, Huang and Sauer 2010). Subsequently, mechanistic target of rapamycin (mTOR) complex 1 (mTORC1) becomes stimulated, therby positively governing Th cell differentiation via downstream signaling factors such as S6K (Kurebayashi, Nagai et al. 2012). PI3K-Akt-mTORC1 signaling also induces expression of hypoxia-inducible factor-1α (HIF-1α) downstream of mTORC1 (Shi, Wang et al. 2011, Ikejiri, Nagai et al. 2012) and in turn, HIF1-α positively regulates Th17 cell differentiation by enhancing Th17-related genes such as RORγt and the glycolytic activity needed to expand T cells (Dang, Barbi et al. 2011, Shi, Wang et al. 2011). In addition, mTORC1 regulates the phosphorylation of STAT proteins during Th1 and Th17 cell differentiation (Delgoffe, Pollizzi et al. 2011). Moreover, deletion of *Rheb*, resulting in dysfunction of mTORC1, selectively reduces the tyrosine phosphorylation of STAT3 and STAT4, which are downstream signaling molecules of IL-6 and IL-12, respectively (Delgoffe, Pollizzi et al. 2011).

Inositol polyphosphate multikinase (IPMK) is a pleiotropic enzyme essential for the production of inositol polyphosphates, such as IP_5_ and IP_6_. With broad substrate specificity, IPMK is also known to display PI3K activity, which forms the second messenger PIP_3_ by phosphorylating PIP_2_ (Resnick, Snowman et al. 2005, Maag, Maxwell et al. 2011). IPMK deletion in mouse embryonic fibroblasts (MEFs) was shown to decrease cellular PIP_3_ levels as well as significantly reduce PIP_3_-dependent Akt phosphorylation in response to growth factors, suggesting IPMK is a physiological PI3K (Maag, Maxwell et al. 2011). Recently, it is revealed that single-nucleotide polymorphisms (SNPs) in IPMK are associated with immune-mediated diseases such as Crohn’s disease (Yokoyama, Wang et al. 2016). Moreover, IPMK coordinates the activity of various biological events in different cell types. In myeloid cells, IPMK was recently shown to control Toll-like receptor signaling and in B cells this kinase is necessary for full activation of Btk signaling (Kim, Beon et al. 2017, Kim, Kim et al. 2019). Accordingly, we asked whether IPMK also plays a critical role in T cell-mediated immune responses. Here, we demonstrate that IPMK plays a key role in Th1- and Th17-dependent immune reactions by controlling Akt-mTOR signaling. Conditional deletion of IPMK in murine CD4^+^ T cells blunts immune homeostasis dependent on Th1 and Th17, but not Th2 cells, rendering mice more susceptible to *Leishmania major* infection and less susceptible to experimental autoimmune encephalomyelitis (EAE). Further, we show that IPMK deletion impairs Th1 and Th17 cell differentiation, which is associated with reduced production of PIP_3_ leading to marked reduction of Akt and its downstream signaling events during T cell activation and differentiation. Thus, our results identify IPMK as a physiological PI3K of Th1 and Th17 cells.

## Results

### Increased expression of IPMK by Th1 and Th17 cells

To address whether IPMK is expressed in effector CD4^+^ T cells, we sorted naive CD4^+^ CD62L^+^CD25^−^CD44^−^ T cells from wild type (WT) mice by fluorescence-activated cell sorting (FACS) and differentiated them into Th1, Th2, and Th17 cells. While Th2 cells modestly express IPMK, IPMK expression is highly upregulated in Th1 and Th17 cells (***Figure supplement 1***). Because IPMK is highly expressed in Th1 and Th17 effector cells, we generated CD4^+^ T cell-specific *Ipmk*-deficient mice by crossing *Ipmk*-floxed mice (*Ipmk*^f/f^) with *Cd4*-Cre mice (***Figure supplement 2A***) to investigate the function of IPMK in CD4^+^ T cell-mediated immunity. We confirmed that the expression of IPMK is absent in CD4^+^ T cells in CD4-specific *Ipmk*-deficient (*Ipmk*^ΔCD4^) mice (***Figure supplement 2B***). To investigate the effects of IPMK on T cell development, we analyzed the phenotypes of T cells in the thymus, peripheral lymph node (pLN), and spleen. Compared with WT control (*Ipmk*^f/f^), *Ipmk*^ΔCD4^ mice showed a normal proportion of double positive (CD4^+^CD8^+^) T cells in the thymus, and CD4^+^ or CD8^+^ single positive T cells in the thymus, pLN, and spleen (***Figure supplement 3A, B***). Further, the proportion of naïve (CD62L^hi^CD44^lo^) and effector/memory (CD62L^lo^CD44^hi^) CD4^+^ and CD8^+^ T cells in pLN and spleen was similar between *Ipmk*^f/f^ and *Ipmk*^ΔCD4^ mice (***Figure supplement 3C, D***). Thus, development of T cells is normal in CD4^+^ T cell-specific *Ipmk* deletion.

### Reduced Th1 immunity in the absence of IPMK in CD4^+^ T cells

Because IPMK is highly expressed in Th1 cells (***Figure supplement 1***), we next investigated the importance of IPMK in Th1-mediated immune responses. For this, we infected *Ipmk*^f/f^ and *Ipmk*^ΔCD4^ mice with *L. major*, an intramacrophage parasite controlled by type 1 immunity in the host (Sacks and Noben-Trauth 2002). *L. major* promastigotes were inoculated into the hind footpads and progression of lesion swelling was measured for 7 weeks after infection (***Figure 1A***). WT BALB/c mice were used for infection control. Compared with *Ipmk*^f/f^ mice, *Ipmk*^ΔCD4^ mice showed significantly increased footpad thickness, which indicates decreased Th1-mediated immune responses. On the other hand, BALB/c mice were more susceptible, as this strain of mice predominantly mounts Th2-mediated immune responses (***Figure 1B***). Because IFN-γ-production by CD4^+^ T cells is critical for the control of *L. major* (Scott, Natovitz et al. 1988), we performed intracellular cytokine staining on cells from spleens and popliteal lymph nodes (pLNs) 7 weeks after infection. The IFN-γ^+^ Th1 population was significantly reduced in the spleen and pLN of *L. major*-infected *Ipmk*^ΔCD4^ mice (***Figure 1C and D***). Thus, these data indicate IPMK deficiency in CD4^+^ T cells dampens the protective Th1 immunity against *L. major* infection decreasing parasite clearance. This emphasizes the importance of IPMK in Th1 immunity.

**Figure 1.**
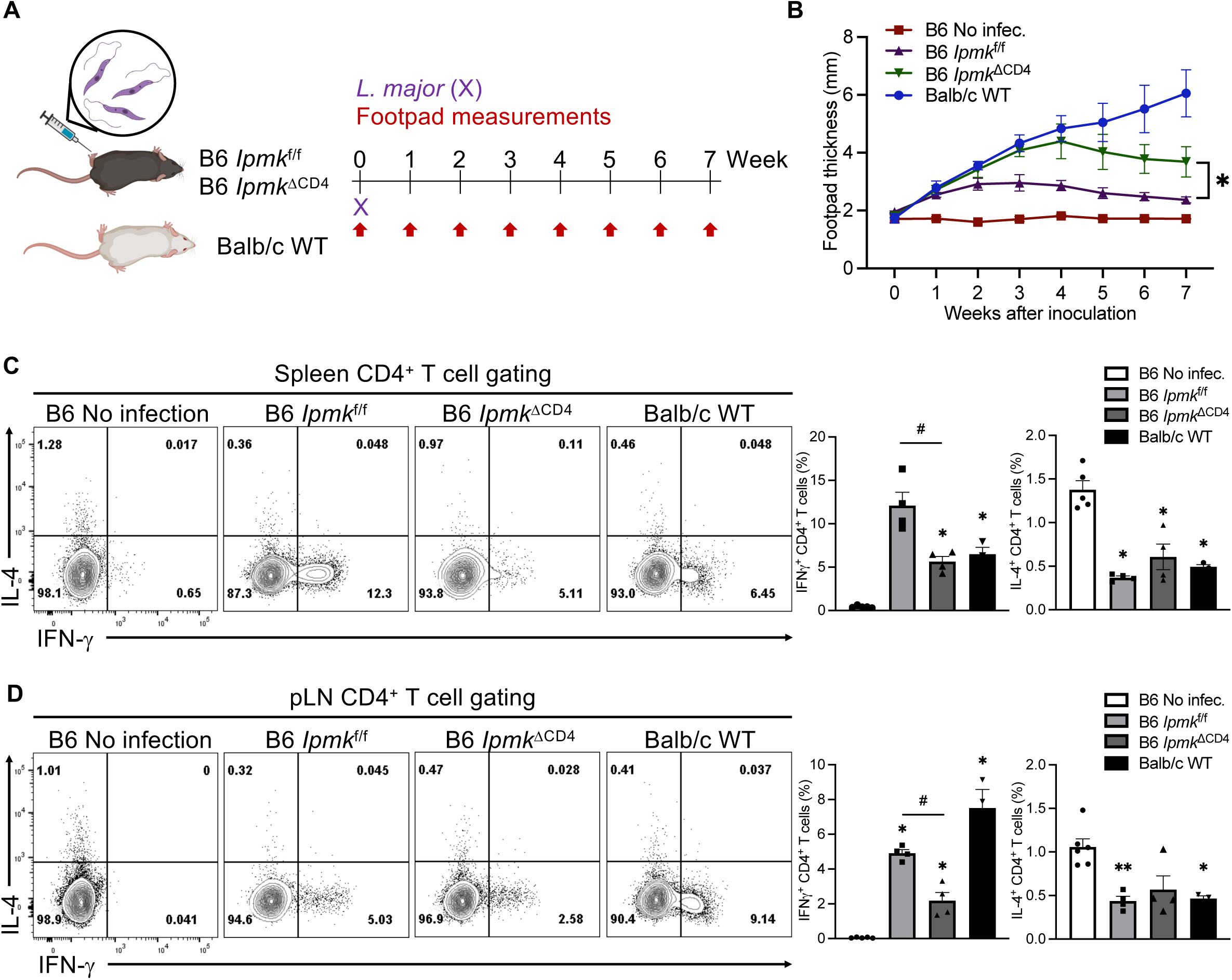
CD4^+^ T cell-specific IPMK deficiency shows reduced Th1-mediated immune responses against infection with *L. major*. Mice were infected with 1 x 10^6^ *L. major* promastigotes in the left hind footpad. (**A**) Schematic diagram of the experimental protocol. (**B**) The footpad swelling was monitored weekly and footpad lesions were measured in no infection (▪), and *Ipmk*^f/f^ (▴), *Ipmk*^ΔCD4^ (▾) in C57BL/6, and Balb/c (●) mice with infection. (**C**) Flow cytometric analyses were performed 7 weeks after infection from spleens. Representative flow cytometric plots and quantification of IFN-*γ*^+^ and IL-4^+^ in CD4^+^ T cells. (**D**) Flow cytometric analyses were performed 7 weeks after infection from popliteal lymph nodes (pLNs). Representative flow cytometric plots and quantification of IFN-*γ*^+^ and IL-4^+^ in CD4^+^ T cells. Data are presented as the mean ± SEM; *\*p* < 0.05 and *\*\*p* < 0.01 compared with the no infection group; by the Mann-Whitney test. ^#^*p* < 0.05 compared with the *Ipmk*^f/f^ group; by the Mann-Whitney test.

### Decreased Th17 cell function due to IPMK deficiency

Next, we investigated the role of IPMK in Th17 cell differentiation and function. For this, we utilized an EAE model in which Th17 cells are known to be critically involved in disease pathogenesis (***Figure 2A***). Following immunization with neuroantigen MOG_35-55_ peptide, *Ipmk*^ΔCD4^ mice showed attenuated disease severity compared with *Ipmk*^f/f^ mice (***Figure 2B***). Histological analysis showed decreased accumulation of infiltrating immune cells and resultant demyelination in the spinal cord of *Ipmk*^ΔCD4^ mice compared with that of *Ipmk*^f/f^ mice (***Figure 2C***). In addition, infiltration of inflammatory mononuclear cells, including CD4^+^ T cells, CD8^+^ T cells, neutrophils, macrophages and B cells, into the central nervous system (CNS) was decreased in *Ipmk*^ΔCD4^ mice compared with that of *Ipmk*^f/f^ mice (***Figure 2D and E***). Indeed, the immunological analysis revealed that there were fewer CNS-infiltrating IL-17A^+^ and IFN-γ^+^ CD4^+^ T cells in *Ipmk*^ΔCD4^ mice than in *Ipmk*^f/f^ mice (***Figure 2F***). These findings indicate that IPMK is required for the function of Th17 cells to mediate autoimmune pathology.

**Figure 2.**
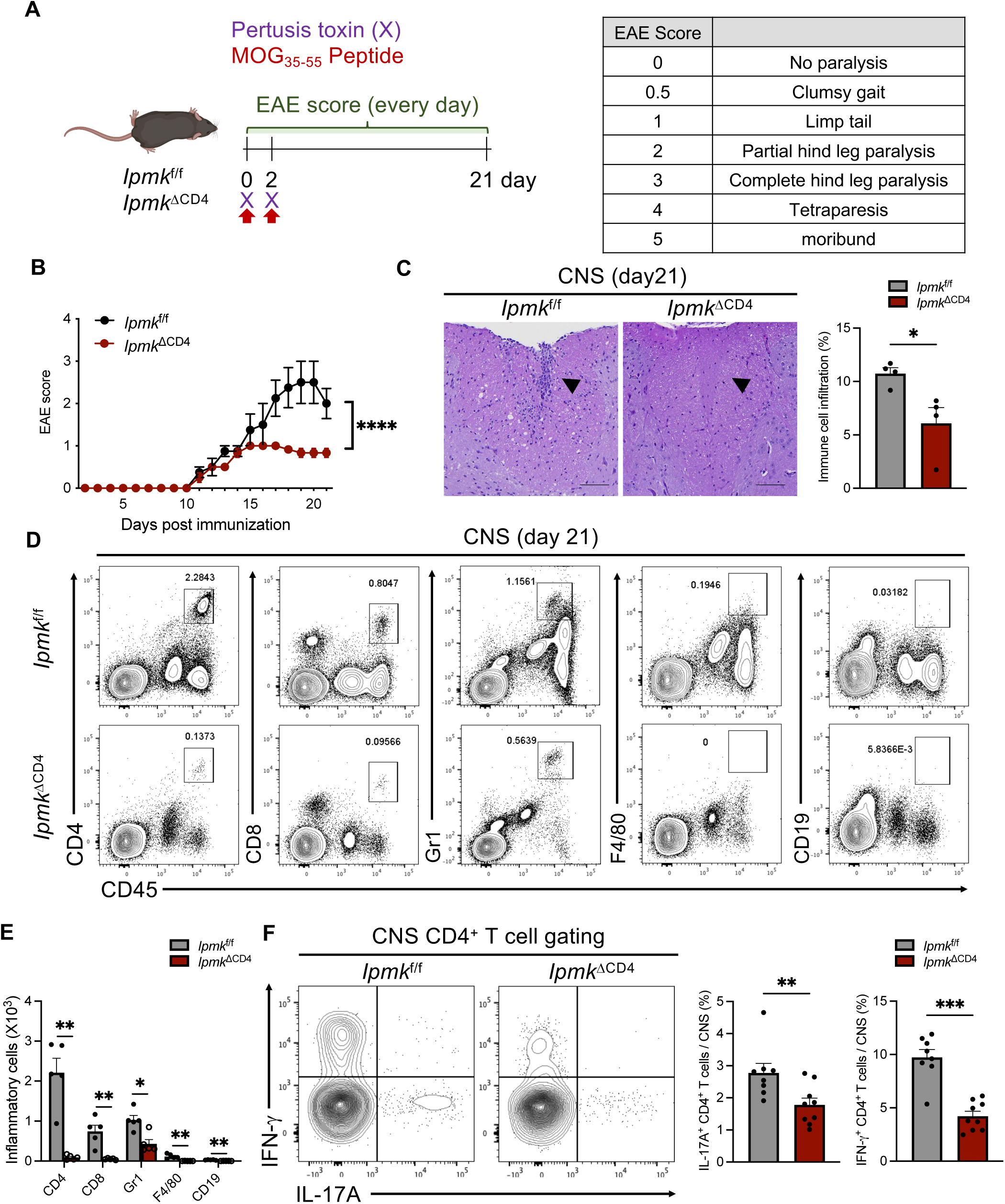
*Ipmk*^ΔCD4^ mice protected from EAE. (**A**) Schematic diagram of the experimental protocol. (**B**) Mean clinical score for EAE in *Ipmk*^f/f^ or *Ipmk*^ΔCD4^ mice induced by immunization with MOG_35-55_. (**C**) Hematoxylin and eosin (H&E) staining from *Ipmk*^f/f^ or *Ipmk*^ΔCD4^ mice harvested at the peak of disease. Scale bar, 100 mm. (**D**) immune cell infiltration in the brains of MOG_35-55_-immunized *Ipmk*^f/f^ or *Ipmk*^ΔCD4^ mice harvested at the peak of disease analyzed by flow cytometry with antibodies to the indicated proteins. (**E**) Absolute numbers of mononuclear CNS-infiltrating cells in harvested brains and spinal cords, determined at the peak of disease by staining with antibodies to the indicated cell markers and subsequent flow cytometry. (**F**) Flow cytometry of IL-17A or IFN-γ production of spinal cord from *Ipmk*^f/f^ or *Ipmk*^ΔCD4^ mice. (n ≥ 8 per each group) Each figure shows representative and compiling data. Data are presented as the mean ± SEM; *\*\*p* < 0.01, *\*\*\*p* < 0.001 and *\*\*\*\*p* <0.0001 compared with the *Ipmk*^f/f^ group; by the Mann-Whitney test.

Next, to test the involvement of IPMK in Th2-mediated immune responses, we induced allergic asthma and analyzed the disease phenotypes and Th2 responses. For this, *Aspergillus oryzae* protease (AP) mixed with chicken egg ovalbumin (OVA; APO) allergen (Kim, Lee et al. 2017, Kwon, Kim et al. 2017, Kim, Kim and Lee 2018) was administrated and asthma phenotypes including airway hyperresponsiveness (AHR), recruitment of eosinophils, goblet cell metaplasia, and mucus hyperproduction, which are mediated by type 2 cytokines IL-4, IL-5, and IL-13, were examined (Lambrecht and Hammad 2015, Walker and McKenzie 2018, Leon and Ballesteros-Tato 2021). APO allergen was intranasally administrated into *Ipmk*^f/f^ and *Ipmk*^ΔCD4^ mice five times and allergic asthma symptoms were analyzed within 16 h of the final challenge (***Figure supplement 4A***). Similar to WT control, *Ipmk*^ΔCD4^ mice also showed enhanced symptoms of allergic asthma including AHR, eosinophilic airway inflammation, mucus hyper-production and periodic acid-Schiff (PAS) positive goblet cells, and increased secretion of type 2 cytokines (***Figure supplement 4B-I***). Thus, these data indicate that IPMK is dispensable for function of Th2 cells, as Th2-mediated immune responses were not affected. Collectively, IPMK-mediated signaling is important for the function of Th1 and Th17 cells, but not of Th2 cells.

### IPMK is required for Th17 cell generation *in vitro*

Next, we investigated the effect of IPMK deficiency on *in vitro* T cell differentiation using naive CD4^+^ T cells from *Ipmk*^f/f^ or *Ipmk*^ΔCD4^. Particularly, we focused on Th17 cells since they exhibit the most robust expression of IPMK (***Figure supplement 1***) and display severe *in vivo* defects in immune responses due to deficiency of IPMK (***Figure 2***). Naive *Ipmk*^ΔCD4^ CD4^+^ T cells displayed significant impairment in their differentiation into Th17 cells as shown by a marked decrease in the IL-17A^+^ cell population compared to the *Ipmk*^f/f^ CD4^+^ T cell population (***Figure 3A***) and diminished secretion of IL-17A into culture supernatants (***Figure 3B***). In addition, analysis of the proliferation of *Ipmk*^ΔCD4^ CD4^+^ T cells revealed a reduction in overall frequency (***Figure 3C and D***) and each proliferation cycle of IL-17A^+^ T cells (***Figure 3D***) compared to that of *Ipmk*^f/f^ cells, suggesting that the impaired function of IPMK KO Th17 cell is not merely a result of reduced proliferation. We also examined whether the kinetics of IPMK expression is dynamically changed under Th17-polarizing conditions and found substantial upregulation of *Ipmk* mRNA 9 h after activation in the presence of IL-6 and TGF-β, which gradually diminished after 24 h (***Figure 3E***). In addition, expression of *Il17a* was substantially decreased in *Ipmk*^ΔCD4^ T cells after 24 h under Th17 cell differentiation conditions (***Figure 3F***). Collectively, these findings strongly suggest that IPMK is a key factor for controlling the expression of IL-17A and for the generation of functional Th17 cells.

**Figure 3.**
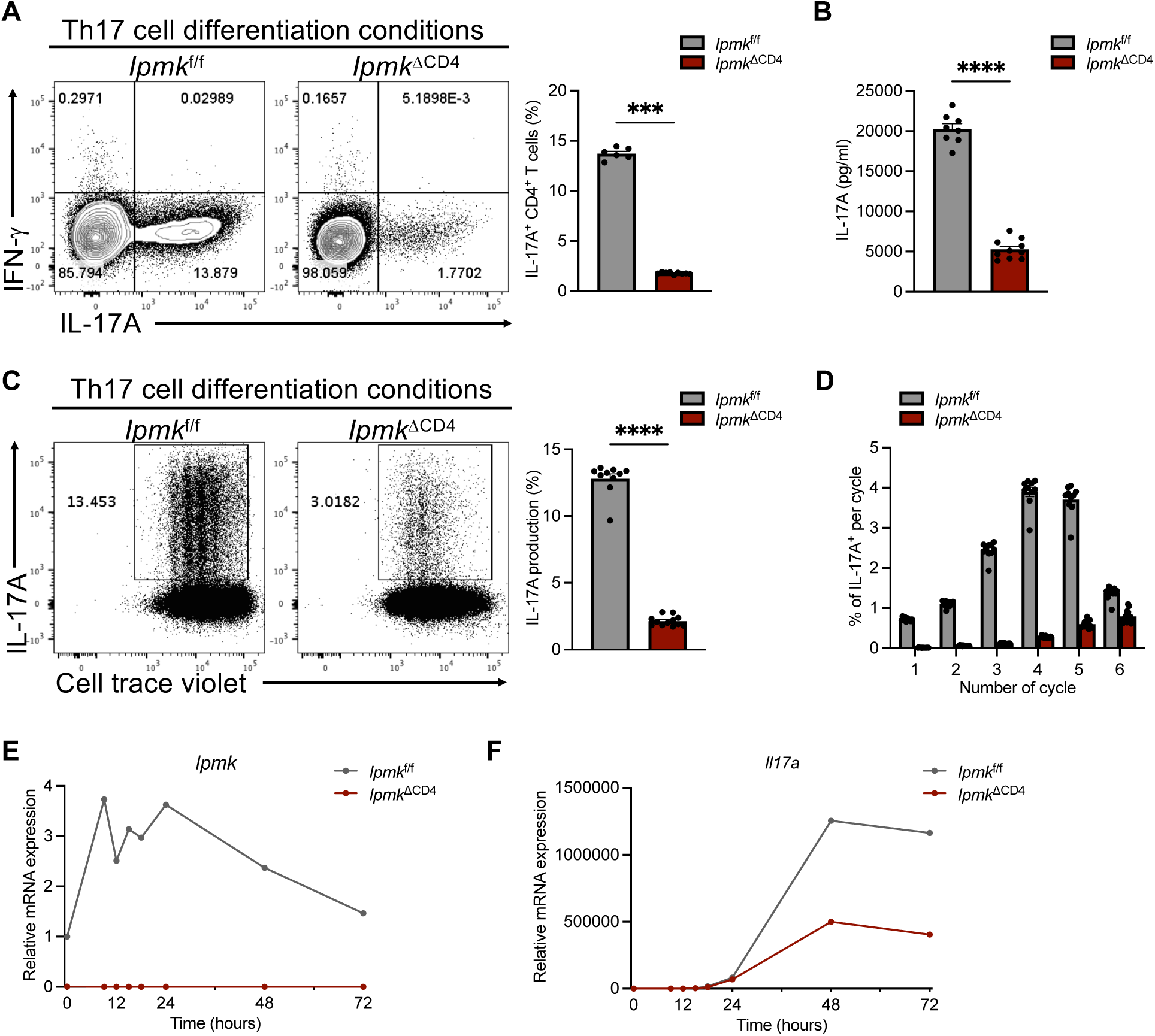
IPMK promotes Th17 cell differentiation. (**A**) Flow cytometry of IL-17A produced by Th17 cells from *Ipmk*^f/f^ or *Ipmk*^ΔCD4^ mice for 3 days *in vitro*. (n ≥ 6 per each group) (**B**) The IL-17A concentration of supernatant from Th17 cells culture conditions. (n ≥ 8 per each group) (**C**) The proliferation of Th17 cells of indicated genotypes for 3 days assessed by cell trace violet dilution assay and flow cytometry. (n ≥ 10 per each group) (**D**) The percentage of IL-17^+^ cells per each cycle from 1 to 8 proliferation as in (**C**) of flow cytometry data. (**E and F**) Quantitative real-time PCR analysis of the time course of the expression of *Ipmk* (**E**) and *Il17a* (**F**) mRNA in naive CD4 T cells polarized into Th17 cells from *Ipmk*^f/f^ or *Ipmk*^ΔCD4^ mice. Each figure shows representative and compiling data. Data are presented as the mean ± SEM; *\*\*\*p* < 0.001 and *\*\*\*\*p* <0.0001 compared with the *Ipmk*^f/f^ group; by the Mann-Whitney test.

### IPMK globally regulates the transcriptional program of Th17 cells

Next, we performed genome-wide RNA sequencing (RNA-seq) analysis to assess the alteration of gene expression in IPMK-deficient cells during Th17 cell differentiation. Principal component analysis (PCA) showed that *Ipmk*^ΔCD4^ Th17 cells (*Ipmk*^ΔCD4^_Th17) were distinct from *Ipmk*^f/f^ Th17 cells (*Ipmk*^f/f^_Th17). Furthermore, *Ipmk*^ΔCD4^ Th0 cells (*Ipmk*^ΔCD4^_Th0) cells also differed from *Ipmk*^f/f^ Th0 cells (*Ipmk*^f/f^_Th0) (***Figure 4A***). Gene ontology analysis revealed that the Th cell differentiation pathway was modulated in *Ipmk*^ΔCD4^ CD4 T cells differentiated into Th17 cells (***Figure 4B***). We next analyzed how IPMK deficiency affects the gene expression of Th17 cells and found that expression of *Rorc, Il17a, Il17f, Il1r2, Tcf7* and *Lef1,* genes related to Th17 transcriptional programs (Crome, Wang and Levings 2010, Muranski, Borman et al. 2011, Yu, Sharma et al. 2011), was significantly decreased in *Ipmk*^ΔCD4^ Th17 cells compared with *Ipmk*^f/f^ Th17 cells. However, expression of Th17 negative regulators, *Il2ra, Foxp3, or Il10,* was markedly increased in *Ipmk*^ΔCD4^ Th17 cells relative to *Ipmk*^f/f^ Th17 cells (***Figure 4-Figure supplement 5***). The validation of IPMK-related genes by reverse transcription quantitative PCR (RT-qPCR) showed that *Rorc, IL17a,* as well as *Il23r*, which are hallmarks of Th17 cells, were upregulated in WT, but not in IPMK-deficient, Th17 cells (***Figure 4D***). Moreover, we confirmed that the expression of *Foxp3*, a key regulator of Treg cell differentiation, and Treg-related cytokine *Il10* was increased in *Ipmk*^ΔCD4^ naive CD4^+^ T cells stimulated under Th17 polarizing conditions, compared to *Ipmk*^f/f^ cells (***Figure 4D***). In addition to mRNA expression, *Ipmk*^ΔCD4^ T cells displayed less IL-17A, but higher Foxp3, protein expression than *Ipmk*^f/f^ T cells during Th17 cell differentiation (***Figure 4E***). Thus, IPMK is required to encourage Th17 cell-related programs and to restrict those of Treg cells during differentiation.

**Figure 4.**
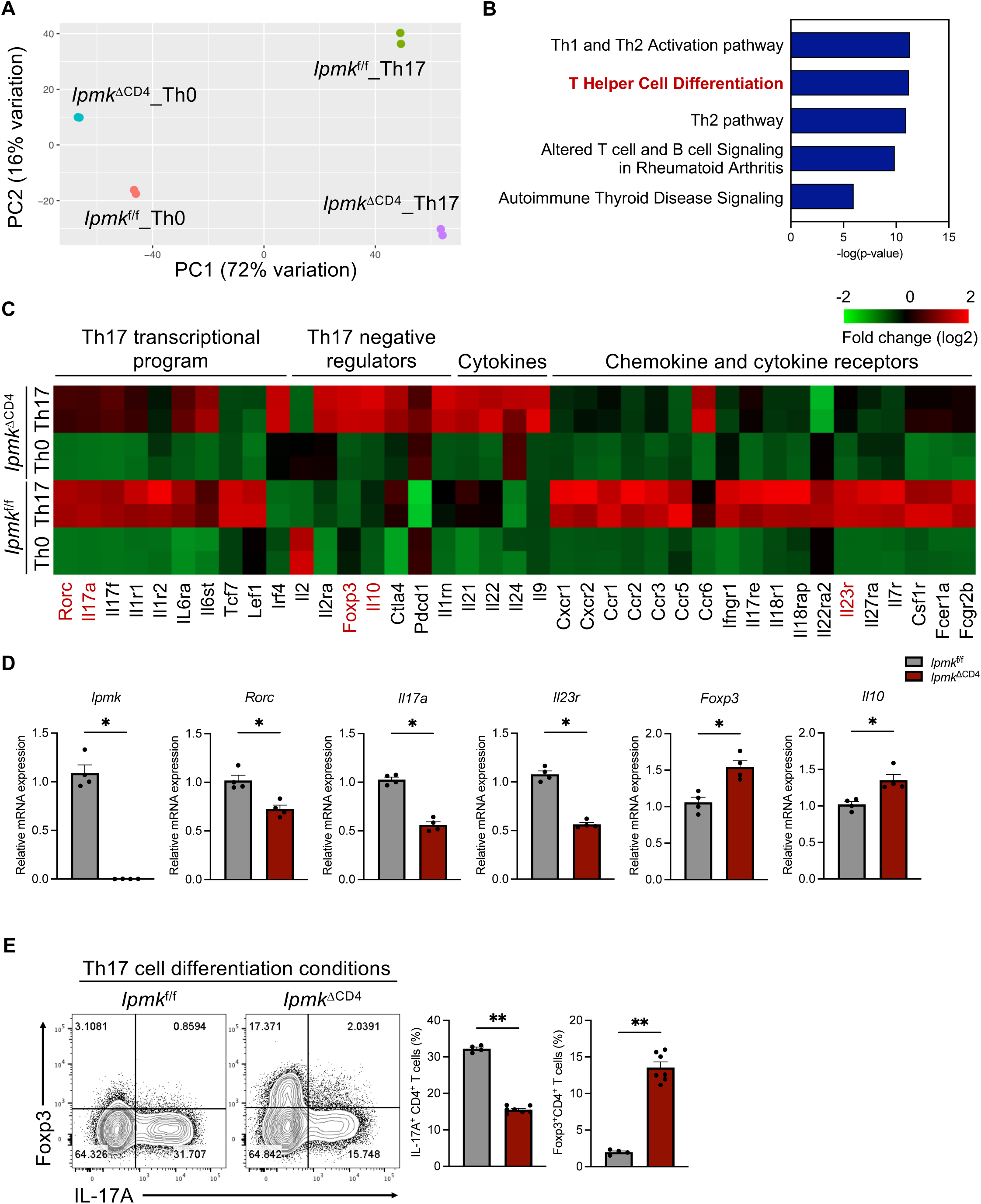
Transcriptionally distinct *Ipmk*^f/f^ or *Ipmk*^ΔCD4^ Th17 cells. (**A**) Principal component analysis of the samples. (n=2) (**B**) Pathway enrichment analysis of genes whose expression is significantly modulated by IPMK in Th17 cells from *Ipmk*^f/f^ or *Ipmk*^ΔCD4^ mice activated under indicated polarizing condition for 3 days. (**C**) Heat map depicting the gene expression (FPKM) in the amount of transcripts and red color significantly and differently expressed between *Ipmk*^f/f^ or *Ipmk*^ΔCD4^ cells stimulated indicated polarizing condition. Data for each experimental group (n=2 per condition) are shown. The color gradient indicates DEG expression as shown. (**D**) Quantitative real-time PCR analysis to detect mRNA levels of Th17 cell-related genes and Treg cell-related genes expressed by CD4^+^ T cells of indicated genotypes, activated under Th17 cell-polarizing condition for 3 days. (n=4) (**E**) Flow cytometry of IL-17A or Foxp3 expressed by nonpathogenic Th17 cell condition of *Ipmk*^f/f^ or *Ipmk*^ΔCD4^ cells for 3 days. (n ≥ 4 per each group) Each figure shows representative and compiling data. Data are presented as the mean ± SEM; *\*p* < 0.05 and *\*\*p* < 0.01 compared with the *Ipmk*^f/f^ group; by the Mann-Whitney test.

### IPMK is critical for STAT and Akt-mTOR signaling pathways in Th17 cells

Next, we further investigated the contribution of IPMK to the control of IL-6 mediated Th17 cell differentiation. STAT3 phosphorylation triggered by IL-6 is central to Th17 cell differentiation (Yang, Panopoulos et al. 2007) and was found to be significantly downregulated in *Ipmk*^ΔCD4^ T cells during Th17 cell generation (***Figure 5A***). Phosphorylation of STAT3 Y705 and S727 residues is commonly regulated through Janus kinase 1 (JAK1), JAK2 (Seif, Khoshmirsafa et al. 2017), and mTORC1 during Th17 cell differentiation (Bezzerri, Vella et al. 2016). Accordingly, gene set enrichment analysis (GSEA) revealed that the mTOR signaling pathway was found to be downregulated (***Figure 5B***) and a reduction in phosphorylation of mTOR was seen in *Ipmk*^ΔCD4^ Th17 cells (***Figure 5C***), suggesting a decrease in mTOR activation in the absence of IPMK. To address the association between phosphorylation of STAT3 and mTOR activity, we examined whether Akt and mTOR activation was affected in *Ipmk*^ΔCD4^ T cells. Diminished phosphorylation of Akt on both T308 and S473 residues was observed in *Ipmk*^ΔCD4^ CD4^+^ T cells at early time point of TCR stimulation (***Figure 5D***). Thus, this suggests IPMK is required for activation of Akt and mTOR signaling pathways in CD4^+^ T cells. To further investigate whether IPMK expression in CD4^+^ T cells mediates Akt activation through controlling PIP_3_ levels, we quantified PtdIns(4,5)P_2_ and PtdIns(3,4,5)P_3_ using LC-MS based on derivatization (Clark, Anderson et al. 2011) in naive CD4^+^ T cells. The ratio of PtdIns(3,4,5)P_3_ to PtdIns(4,5)P_2_ was decreased in *Ipmk*^ΔCD4^ naive CD4^+^ T cells stimulated with anti-CD3 and anti-CD28 antibodies (***Figure 5E***). This data indicates that the catalytic function of IPMK as a PI3K is necessary for the production of PtdIns(3,4,5)P_3_ from PtdIns(4,5)P_2_ in CD4^+^ T cells, which subsequently stimulate the Akt-mTOR signaling axis to drive Th17 cell differentiation.

**Figure 5.**
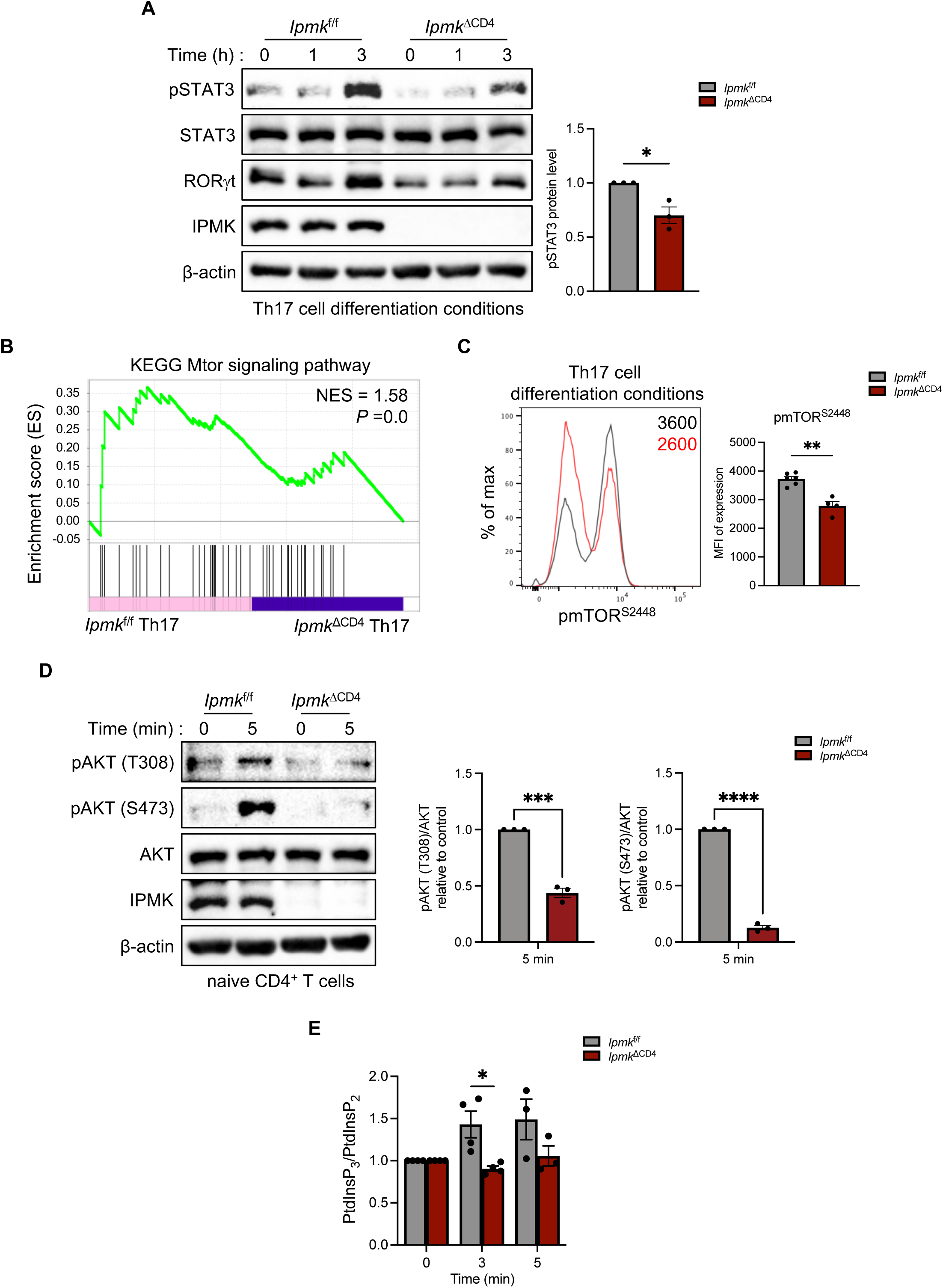
Defective activation of the Akt-mTOR pathway in *Ipmk*^ΔCD4^ CD4^+^ T cells. (**A**) Immunoblotting of STAT3 phosphorylated at Y705 or total STAT3 in lysates of naive CD4^+^ T cells polarized for 3 days under Th17 conditions from *Ipmk*^f/f^ or *Ipmk*^ΔCD4^ then restimulated for carious times with IL-6 and TGF-β. Unpaired Student’s *t* test was used for statistical analysis; *\*p* < 0.05 compared with the *Ipmk*^f/f^ group. (**B**) GSEA-based KEGG-enrichment plots for downregulated genes in Mtor signaling pathway of *Ipmk*^ΔCD4^ Th17 cells. (**C**) Sorted naïve CD4 T cells were differentiated under Th17 condition and evaluated for levels of pmTOR (S2448). (**D**) Immunoblotting of AKT phosphorylated at T308, S473 and total AKT in lysates of naive CD4^+^ T cells stimulated with anti-CD3, anti-CD28, and hamster IgG antibodies. Unpaired Student’s *t* test was used for statistical analysis; *\*\*\*p* < 0.001; *\*\*\*\*p* < 0.0001 compared with the *Ipmk*^f/f^ group. (**E**) Measuring the ratio of PtdInsP_3_ to PtdInsP_2_ in Naive *Ipmk*^f/f^ or *Ipmk*^ΔCD4^ CD4^+^ T cells, which were stimulated with anti-CD3, anti-CD28, and hamster IgG antibodies. PtdInsP_3_ and PtdInsP_2_ levels was quantified by LC-MS/MS (technical replicates n=4). *\*p* < 0.05 compared with the *Ipmk*^f/f^ group; by the Mann-Whitney test. Each figure shows representative and compiling data. Data are presented as the mean ± SEM.

### IPMK-deficient CD4^+^ T cells exhibit an altered metabolic profile

In addition to STAT3 activation, mTOR has been shown to play an important role in the regulation of T cell metabolism through activation of glycolysis-related genes (Linke, Fritsch et al. 2017). Notably, T cell differentiation requires the activity of mTOR-dependent pathways, and in particular, differentiation of Th1 and Th17 cells is induced through mTOR-dependent glycolysis. Therefore, we performed a mitochondrial stress test to assess mitochondrial respiration via the oxygen consumption rate (OCR) using the Seahorse extracellular flux analyzer. *Ipmk*^ΔCD4^ T cells exhibited lower basal OCR, maximal mitochondrial respiration, spare respiratory capacity, and ATP production than did *Ipmk*^f/f^ CD4^+^ T cells (***Figure 6A and B***). We also assessed energy production through glycolysis using the XF glycolysis stress test. The basal extracellular acidification rate (ECAR), which correlates with lactate production by glycolysis, was reduced in *Ipmk*^ΔCD4^ T cells but not in *Ipmk*^f/f^ T cells. However, there were no differences in glycolytic capacity or glycolytic reserves between the two groups after injection of an ATP synthase inhibitor, oligomycin (***Figure 6C and D***). Consistently, we found that *Ipmk*^ΔCD4^ T cells exhibited a reduced oxygen consumption rate (OCR) and glycolytic energy production following the addition of oligomycin. Collectively, these results suggest that IPMK deficiency leads to impaired mitochondrial energy production and glycolysis in CD4^+^ T cells in a manner dependent on the STAT3-mTOR pathway.

**Figure 6.**
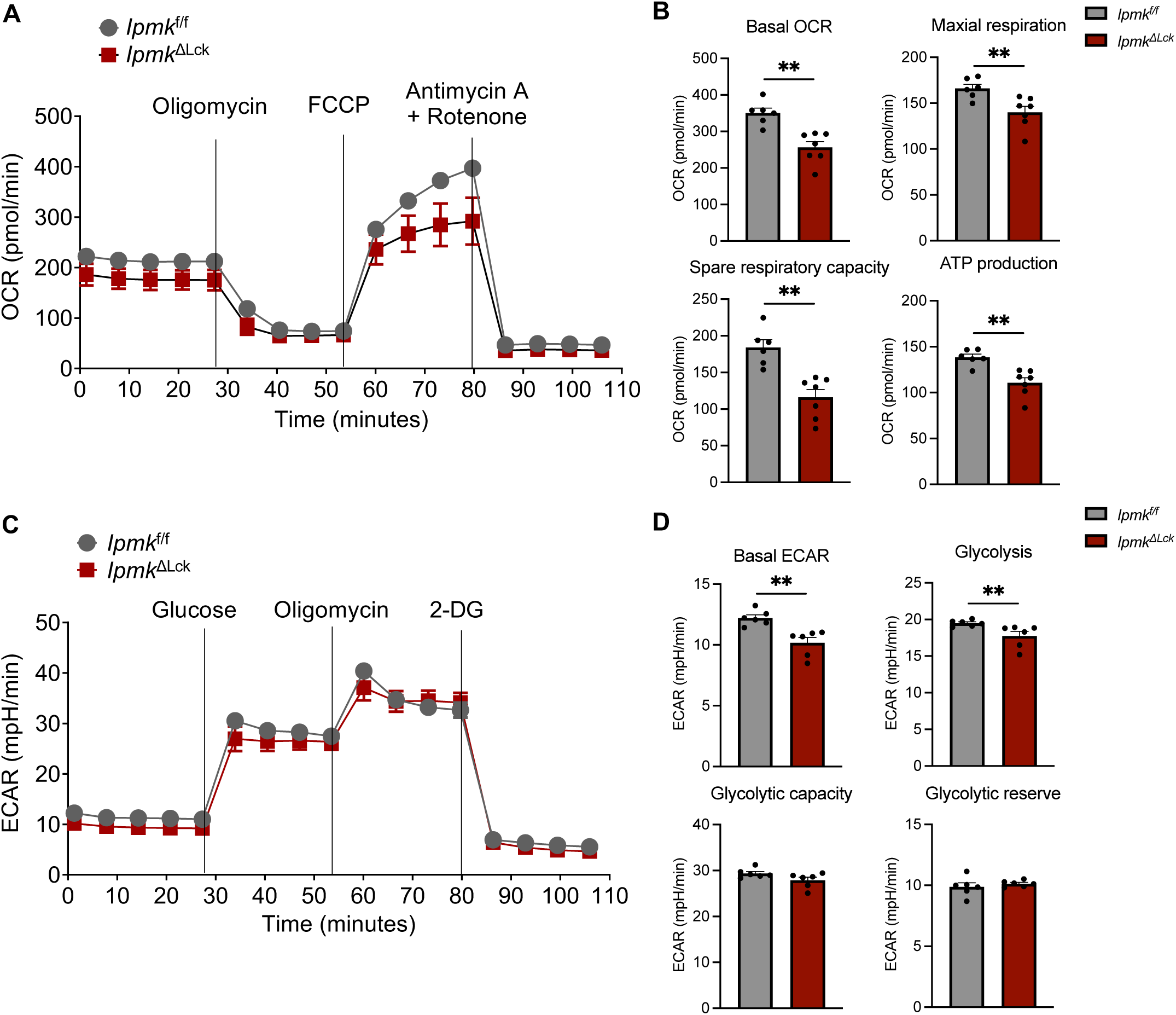
Reduced aerobic glycolysis and mitochondrial oxidative respiration with IPMK in CD4^+^ T cells. (**A-D**) CD4 cells from *Ipmk*^f/f^ or *Ipmk*^ΔLck^ were activated with dynabead for 72 hours. OCR (**A**) was measured by Seahorse XFe Analyzer and calculated basal OCR, maximal respiration, spare respiratory capacity and, ATP production (**B**). ECAR (**C**) was measured by Seahorse XFe Analyzer and calculated basal ECAR, glycolysis, glycolytic capacity, and glycolytic reserve (**D**). Each figure shows representative and compiling data. Data are presented as the mean ± SEM; *\*\*p* < 0.01 compared with the *Ipmk*^f/f^ group; by the Mann-Whitney test.

## Discussion

In this study, we demonstrate that IPMK expression is elevated under Th1 and Th17 polarizing conditions, but not under Th2-polarizing conditions. To evaluate the role of IPMK in Th1 and Th17 cell functions, we used CD4^+^ T cell-specific genetic deletion of IPMK and found that the Th1 response was diminished as exhibited by enhanced susceptibility to *L. major* infection, which requires IFN-γ dependent Th1-mediated responses for clearance. Further, EAE, a Th17-dependent autoimmune disease model, exhibited delayed onset and a significantly attenuated Th17 response in *Ipmk*^ΔCD4^ mice. Moreover, IL-17A production was dramatically reduced in IPMK-deficient CD4^+^ T cells compared with WT CD4^+^ T cells under Th17 polarizing conditions. Mechanistically, we showed that IPMK participates in cytokine signaling of STAT activation via the PI3K-Akt-mTOR pathway, suggesting IPMK is a critical regulator of Th1/Th17 cell differentiation (***Figure 7***).

**Figure 7.**
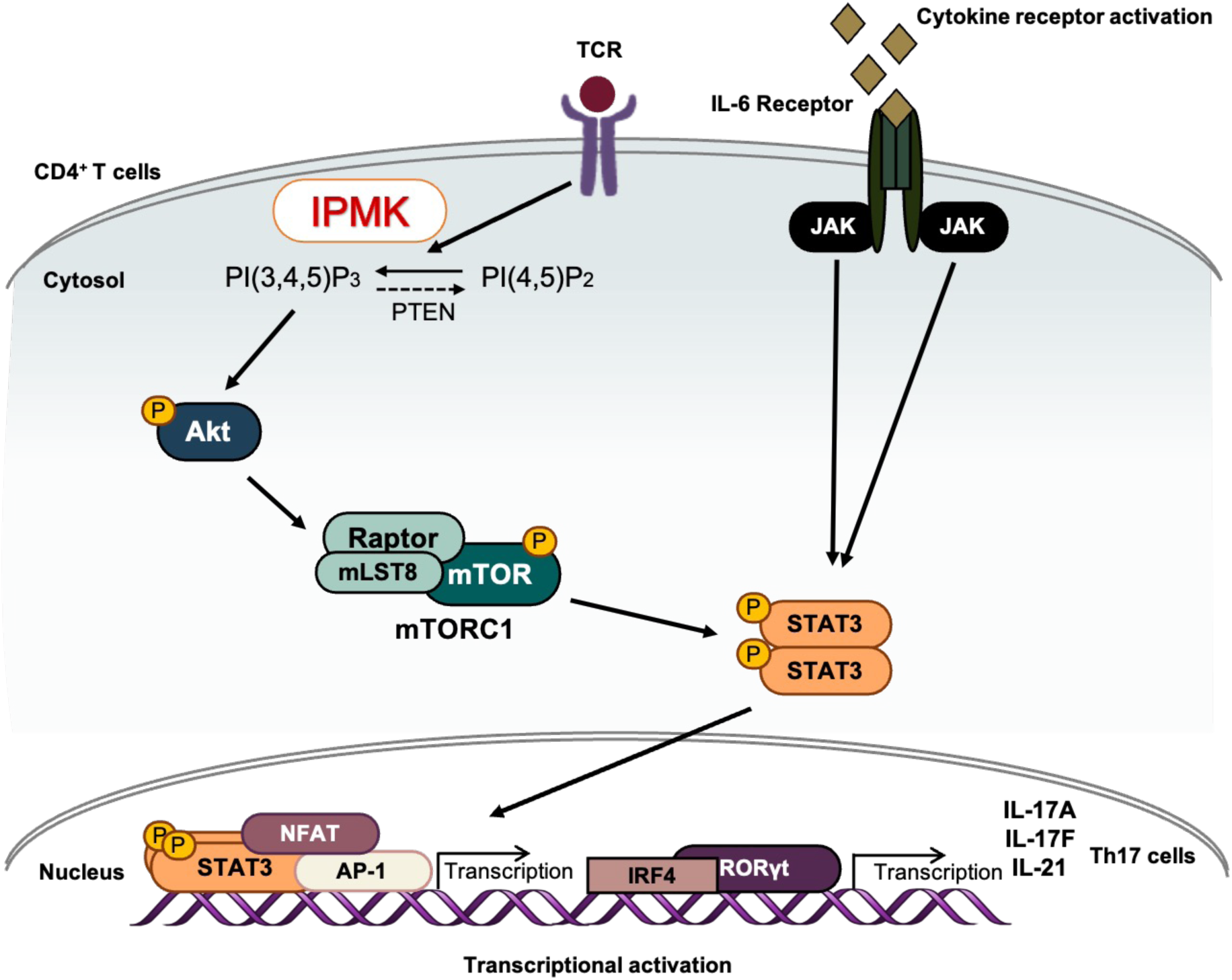
A model deciphering action of IPMK in CD4 T cells under Th17 differentiation condition. Graphic abstract showing how CD4 IPMK signaling regulates Th17 cell differentiation. IPMK activation by IL-6 and TCR signaling causes Akt-mTORC1 and STAT3 phosphorylation. These changes collectively induce Th17 differentiation.

IPMK is known to modulate various biological processes by working enzymatically to catalyze inositol polyphosphate biosynthesis (Saiardi, Erdjument-Bromage et al. 1999, Odom, Stahlberg et al. 2000) and production of PIP_3_ (Maag, Maxwell et al. 2011), and non-catalytically by regulating key signaling factors (Kim, Kim et al. 2011, Kim, Beon et al. 2017). For T cells it is reported that IP_4_ generated by ItpkB binds a single ITK subunit causing its recruitment to the plasma membrane and resulting in amplification of TCR signaling, triggering thymocyte positive selection (Huang, Grasis et al. 2007). However, there are no reports as to whether IPMK has a role in CD4^+^ T cell differentiation and function. Therefore, we focused on IPMK in CD4^+^ Th cell differentiation, postulating that IPMK may be particularly important in both Th1 and Th17 responses and possibly Th1 or Th17 cell differentiation.

Upon activation, naive CD4^+^ T cells differentiate into effector T cells including Th1, Th2 and Th17 cells which differently regulate immunity against various pathogens. These effector T cells are differentially induced by distinct environmental cytokines, which signal through STATs or ubiquitous and other inducible transcription factors (Zhu, Yamane and Paul 2010). Interestingly, both IPMK protein and transcripts are significantly increased in Th17 cells. In addition, *Ipmk*^ΔCD4^ T cells cannot be differentiated into Th17 cells, showing attenuated production of IL17A relative to that of *Ipmk*^f/f^ Th17 cells. Differentiation of Th17 cells requires a variety of transcription factors such as STAT3, RORγt, Ahr, BATF, IRF4, etc (Dong 2011, Ciofani, Madar et al. 2012, Yosef, Shalek et al. 2013). Genome-wide sequencing analyses revealed that under Th17-polarizing conditions, *Ipmk*^ΔCD4^ T cells show decreased *Il17a, Rorc,* and *Irf4* transcription. Surprisingly, expression of Foxp3, a master transcription factor for Treg cells (Li, Li et al. 2015), was the most upregulated gene in *Ipmk*^ΔCD4^ Th17 cells. Foxp3^+^ Treg cells are critical for maintenance of immune tolerance and control of immune responses toward pathogens. Interestingly, Th17 and Treg cells are closely related and reciprocally regulated as they share TGF-β cytokine signaling. Therefore, our findings propose IPMK as the key factor to balance Th17 and Treg cell development critical for host protective immunity.

During T cell activation, ligation of receptors by costimulatory molecules induces PI3K activation and consequently activates mTOR, which induces the differentiation of effector T cells. The two mTOR complexes, mTORC1 and mTORC2, have disparate effects on effector T cell differentiation (Delgoffe, Kole et al. 2009, Chi 2012). *Rheb*-deficient CD4^+^ T cells exhibit selectively abrogated tyrosine phosphorylation of STAT3 and STAT4, which is a result of increased suppressor of cytokine signaling 3 (SOCS3) protein expression (Delgoffe, Pollizzi et al. 2011). The mTOR-deficient T cells are unable to activate lineage-specific STAT proteins and major transcription factors. Rheb protein deficiency and the consequent loss of mTORC1 activity leads to the impaired differentiation of Th1 and Th17 cells. Thus, this suggests that mTORC1 mediates the mTOR-dependent differentiation of Th1 and Th17 cells. The findings in the present study suggest that through its PI3K activity IPMK is a key factor mediating the full activation of Akt-mTOR signaling. Specifically, this is shown by the reduced PIP_3_ generation in *Ipmk*^ΔCD4^ T cells. In activated T cells, class I PI3K p110δ has been shown to be a primary enzyme involved in the generation of PIP_3_, thus governing proliferation, cytokine production, and differentiation into Th subsets (Okkenhaug, Bilancio et al. 2002, Okkenhaug, Patton et al. 2006, Sinclair, Finlay et al. 2008, Huang and Sauer 2010). Although IPMK exhibits robust PI3K activity *in vitro* and IPMK-depletion in cell lines (e.g. heterologous MEFs and cancer cells) reduces growth factor-stimulated PIP_3_ synthesis accompanied by hypoactivation of Akt-mTOR signaling, it was challenging to elucidate the appropriate physiological context in which IPMK’s PI3K activity fulfills a central role in Akt-mTOR activation. In B cells, IPMK’s catalytic action in water-soluble inositol polyphosphate biosynthesis is critical for LPS-triggered B cell receptor signaling via IP_6_-dependent Btk activation (Kim, Kim et al. 2019). In macrophages, IPMK seems to non-catalytically function as a scaffold to stabilize TRAF6 and transmit TLR4 signals (Kim, Beon et al. 2017). Recently, conditional deletion of IPMK in Treg cells revealed its essential role for the regulatory function and differentiation of Treg cells into effector Treg cells. Mechanistically, IPMK’s catalytic activity as an IP kinase was found to promote InsP_3_-mediated store-operated Ca^2+^ entry (SOCE) via IP_4_ [Ins(1,3,4,5)P_4_] production in Treg cells (Min, Kim et al. 2022). In contrast, our results in the present study revealed that Th cells depend on the lipid PI3-kinase function of IPMK to regulate activation of mTOR signaling, which directly modulates STAT3 activation and governs Th1 and Th17 cell differentiation. Future studies are needed to define the signaling mechanisms by which IPMK in Th cells cooperates with other classical PI3Ks, such as P110δ, to fully mediate T cell activation, differentiation, and homeostasis. The recent discovery of single-nucleotide polymorphisms (SNPs) in IPMK in immune-mediated diseases (e.g. rheumatoid arthritis, psoriasis, and Crohn’s disease) further highlights the possible contribution of IPMK to the onset and progress of Th cell-mediated human diseases (Yokoyama, Wang et al. 2016).

In summary, our study explores the role of IPMK in proinflammatory CD4^+^ T cell function. Moreover, deletion of IPMK in CD4^+^ T cells attenuates resistance to *L. major* infection and the severity of EAE by impairing Th1 and Th17 cell differentiation. Given the vital importance of Th1 and Th17 cells in T cell immunity, we suggest that regulation of IPMK in CD4^+^ Th cells could be beneficial for the treatment of infections as well as autoimmune disorders.

## Materials and Methods

### Mice

*Cd4*-Cre, *Ipmk*^flox/flox^ and WT mice were on the C57BL/6 background. Mice used for experiments were 6-10 weeks old male and female mice and age-matched. Littermates were used unless stated otherwise. All mice were housed and bred in specific pathogen-free conditions and fed with a standard normal diet *ad libitum* with free access to water. All animal experimental procedures were performed in accordance with guidelines approved by the Korea Advanced Institute of Science and Technology Animal Care and Use Committee (KAIST, KA2013-32 and KA2018-52).

### CD4^+^ T cell differentiation and proliferation *in vitro*

Naive CD4^+^ T cells were isolated from the spleen and pLNs by mouse naive CD4^+^ enrichment kit (Invitrogen) per the manufacturer’s protocols. Followed that, CD4^+^CD25^−^ CD62L^high^CD44^low^ cells were sorted using a FACSAria II (BD Biosciences) equipped with 405 nm, 488 nm, 633 nm lasers and an 85 μm nozzle using a purity mask. Post-sort purity was routinely > 97%. Isolated T cells were plated into plates coated with 10 μg/mL anti-CD3 and 10 μg/mL anti-CD28 antibodies and cultured in RPMI 1640 (Welgene) or IMDM medium (Gibco). TCR stimulation in isolated T cells also progressed with using 20 μg/mL anti-Armenian hamster IgG secondary antibodies for 3 or 5 minutes, which were incubated with 5 μg/mL anti-CD3, 2 μg/mL anti-CD28 antibodies first. Isolated naive CD4^+^ T cells were differentiated to several Th types by different recombinant cytokines and antibodies: Th0 (5 μg/mL anti-IFN-γ, 5 μg/mL anti-IL-4 in RPMI 1640 medium), Th1 (20 ng/mL IL-12, 20 ng/mL IL-2, 5 μg/mL anti-IL-4 in RPMI 1640 medium), Th2 (50 ng/mL IL-4, 10 μg/mL anti-IL-12, 5 μg/mL anti-IFN-γ in RPMI 1640 medium) Th17 (40 ng/mL IL-6, 2 ng/mL TGF-β, 5 μg/mL anti-IFN-γ and 5 μg/mL anti-IL-4 in IMDM medium). To assess proliferation, isolated CD4^+^ T cells were labeled with 5 mM CellTrace Violet Cell proliferation (Life technologies) per the manufacturer’s protocols. Labeled T cells were activated under Th17 differentiation conditions as indicated. The T cell proliferation was assessed 72 h post-activation based on CellTrace Violet dilution by flow cytometry.

### Western blot analysis

Whole-cell lysates were prepared in lysis buffer (1% NP-40, 137 mM NaCl, 20 mM Tris-HCl (pH 8.0), 2 mM EDTA, 10% glycerol, 20 mM NaVO_4_, 10 mM sodium pyrophosphate, 100 mM sodium fluoride, 20 mM PMSF) containing protease inhibitor cocktail (Roche). Whole cells were incubated on ice for 20 min and collected by centrifugation at 17,000g for 15 min. The total protein concentrations were determined by the Bradford protein assay (Bio-Rad), and proteins were boiled at 95 °C for 5 min with SDS sample loading buffer (60 mM Tris-HCl, 25% glycerol, 2% SDS, 14.4 mM β-mercaptoethanol, and 0.1% bromophenol blue). Cell-lysate proteins were electrophoresed on 10% SDS-PAGE gels and separated proteins were transferred onto a nitrocellulose membrane. After blocking with 5% skim milk in Tris-buggered saline (TBST, 0.1% Tween-20) for 1 h at room temperature, membranes were incubated with primary antibodies overnight at 4 °C. Then, the membranes were treated with an HRP-conjugated secondary antibody for 1 h at room temperature. After each step, the membranes were rinsed three times for 10 min with TBST. The HRP signals were developed with Clarity ECL substrate (Bio-Rad) and visualized by ChemiDoc (Bio-Rad) using the Image lab program.

### Flow cytometry

For analysis of surface markers, cells were stained with a combination of the fluroscence-conjugated antibodies, which are listed in ***Supplementary Table 1***, for 20 min at 4 °C in PBS containing 0.25% BSA and 0.02% azide. Dead cells were excluded by LIVE/DEAD Fixable Dead Viability Dye (Invitrogen). For assessment of cytokine production by *in vitro* derived T cells, cells were stimulated with 50 ng/mL phorbol myristate acetate (PMA) and 500 ng/mL Ionomycin with Brefeldin A (Invitrogen; 1:1000) for 3 h then stained for CD4 and a LIVE/DEAD Fixable Dead Viability Dye (Invitrogen). Cells were then fixed and permeabilized (Invitrogen) and stained for IL-17A. For assessment of Foxp3 expression or Il-17A expression, cells were fixed and permeabilized with a Foxp3 staining kit (Invitrogen). Flow cytometry was performed on an LSRII Fortessa (BD Biosciences).

### *Leishmania major* parasites and infectious challenge

*L. major* strain MRHO/SU/59/P/LV39 was cultured and used to infect mice as described (Corry, Reiner et al. 1994, Soong, Xu et al. 1996). Seven weeks following infection, mice were sacrificed and popliteal lymph nodes (pLNs) and spleens were harvested, minced and homogenized using CM for intracellular cytokine assay.

### Intracellular cytokine staining (ICCS)

Single-cell suspensions were re-stimulated PMA (50 ng/mL; Sigma-Aldrich) and ionomycin (500 ng/mL; Sigma-Aldrich) for 5 h at 37 °C in a humidified 5% CO_2_ atmosphere. After stimulation, cells were stained with Fixable Viability dye (Invitrogen), APC-Cy7-conjugated anti-CD8, PE-Cy7-conjugated anti-CD4, and AF700-conjugated anti-CD44 antibodies against the corresponding cell surface proteins. Following the surface staining, samples were fixed and permeabilized using the Intracellular Fixation & Permeabilization Buffer Set from eBioscience, and intracellular IL-4 and IFN-γ were detected by immunostaining with PE-conjugated anti-IL-4 and FITC-conjugated anti-IFN-γ antibodies. Appropriate PE and FITC-conjugated, isotype-matched, irrelevant monoclonal antibodies were used as isotype controls. Flow cytometry was performed using MACSQuant (Miltenyi Biotec, Bergisch Gladbach, Germany) and data were analyzed with FlowJo software (FlowJo, LLC.).

### EAE model

EAE was induced by subcutaneous immunization with 200 μg MOG_35–55_ peptide (Peptron, Daejeon, Korea) emulsified in complete Freund’s adjuvant (Sigma-Aldrich), followed by intravenous injection of 200 ng pertussis toxin (Sigma-Aldrich) on days 0 and 2. To quantify disease severity, scores were assigned daily in a blinded manner on a scale of 0–5 as follows: 0, no paralysis; 0.5, clumsy gait; 1, limp tail; 2, limp tail and partial hind leg paralysis; 3, complete hind leg paralysis; 4, tetraparesis; 5, moribund. Animals were euthanized if scores reached grade 4. To determine CNS infiltrates, cell suspensions from brain and spinal cord were prepared, as described previously (Pino and Cardona 2011).

### Induction of experimental allergic asthma

*A. oryzae* protease (AP) and chicken egg ovalbumin (OVA) were purchased (Sigma-Aldrich) and reconstituted to 1 mg/mL and 0.5 mg/mL using sterile PBS. AP and OVA were mixed to prepare APO allergen at a 1:9 (v/v) ratio immediately before administration. Mice were challenged five times intranasally with 50 μL of APO allergen every 4 days (days 0, 4, 8, 12, and 16). For the intranasal challenge, mice were lightly anesthetized by isoflurane inhalation (Abbott Laboratory). After allergen challenge, AHR, bronchoalveolar lavage (BAL) cytology, BAL glycoprotein assay, and lung histopathology were determined as previously described (Kim, Lee et al. 2017).

### Measurement of airway hyperresponsiveness (AHR)

AHR was measured with a flexiVent system (SciTech, Montreal, Canada). Briefly, 16 h after the final intranasal challenge, mice were anesthetized with pentobarbital (Hanlim Pharma Co., Seoul, Korea) via intraperitoneal injection of 0.15 mL/10 g body weight and intubated with a 20-gauge cannula. After intubation, mice were injected with 0.1 mL/10 g body weight of pancuronium (0.1 mg/mL; Sigma-Aldrich) and ventilated with the flexiVent system. AHR was assessed by administering incremental doses of nebulizing methacholine (0, 1, 3, 9, 27, 81 mg/mL; Sigma-Aldrich) and measuring resistance every 30 seconds. After measuring AHR, BAL fluid (BALF) samples were obtained by washing the lungs with PBS (1 mL, 4 °C) delivered via a tracheal tube.

### Quantification of secreted glycoprotein

Secreted glycoprotein levels in BALF were measured by modified ELISA using jacalin, a glycoprotein-binding lectin. Briefly, a glycoprotein mucin standard derived from porcine stomach (Sigma-Aldrich) and BALF samples were diluted 2-fold serially with PBS, beginning at a 1:100 dilution. 40 μL of each sample was transferred to a flat-bottom ELISA plate (Greiner, Kremsmunster, Austria) and incubated at 4 °C overnight. After washing, the plates were blocked by adding 200 μL of 0.2% I-block (Applied Biosystems) and incubating at 37 °C for 2 h. Plates were washed again, then 40 μL of biotinylated jacalin (Vector Laboratories) diluted in PBS/Tween containing 0.1% BSA (1:1000 dilution) was added and the plates were incubated at room temperature for 30 min. After a final wash, 70 μL of alkaline phosphatase substrate (5 mM *p*-nitrophenyl phosphate substrate in 0.1 M alkaline buffer; Sigma-Aldrich) was added and color was allowed to fully develop. The reaction was terminated by adding 40 μL 0.5 N sodium hydroxide, and optical density was measured at 405 nm using an ELISA plate reader (Marshall Scientific).

### ELISA for cytokine detection

Detection of cytokines in BALF was performed by sandwich ELISA using 96-well plates (Greiner) according to the manufacturer’s instructions (BD Biosciences). Briefly, ELISA plates were pre-coated with 40 μL of 1:500-diluted capture antibody, incubated for 2 h at 37 °C, and then washed and blocked with 200 μL of 0.2% I-block (Applied Biosystems) overnight at 4 °C. After washing, 40 μL of the collected sample was transferred to a 96-well plate and 40 μL of 1:500-diluted detection antibody was added. After incubation for 2 h at 37 °C, plates were washed again and 40 μL of streptavidin-alkaline phosphatase (BD Biosciences) diluted 1:1000 in PBS/Tween/BSA was added and plates were incubated at room temperature for 30 min. After a final wash, 70 μL of alkaline phosphatase substrate (5 M *p*-nitrophenyl phosphate substrate in 0.1 M alkaline buffer; Sigma-Aldrich) was added and plates were incubated at room temperature to allow color development. The reaction was terminated by adding 40 μL of 0.5 N sodium hydroxide, and optical density was measured at 405 nm using an ELISA plate reader (Marshall Scientific).

### Lung tissue histology

For staining with PAS, mouse lungs were inflated and fixed with 4% paraformaldehyde (Sigma-Aldrich) after collection of BALF cells. Lung samples were embedded in paraffin and sectioned (6 μm thick), and then stained with PAS.

### RNA isolation and quantitative RT-PCR

For qRT-PCR analysis, total RNA was extracted from T cells by using RNeasy mini kit (QIAGEN) and reverse-transcribed into cDNA with Superiorscript III reverse transcriptase (Enzynomics, Daejeon, Korea), dNTP set and oligo(dT) primer (Invitrogen) per manufacturer’s protocols. Quantitative PCR (qPCR) was performed using the SYBR Green Master Mix (Roche) and the StepOnePlus Real-Time PCR System (Applied Biosystems). Δ*C*_t_ values were calculated as 2^(reference − gene)^, where the reference was β-actin.

### RNA sequencing

The alignment file was used to perform bulk RNA-sequencing of control or *Ipmk*^ΔCD4^ T cells differentiated into Th17 cells. Briefly, the TopHat software tool was used to map reads. The alignment file was used to assemble transcripts, estimate their abundances, and use cufflinks to detect differential expression of genes or isoforms. The data were further evaluated in the context of canonical signaling using the ingenious pathway analysis program (QIAGEN) to see if *Ipmk*^ΔCD4^ Th17 genes were specifically associated with differentiation. Heatmaps were generated using MeV (http://www.tm4.org). For GSEA, gene set collections from the Molecular Signatures Database 4.0 (http://www.broadinstitute.org/gsea/msigdb/) were used. Sequence data were deposited in the NCBI GEO (accession number GSE203137)

### Cytometric Bead Array (CBA)

Supernatants obtained from the 3 days cultures of differentiated T cells were analyzed for Th cytokines using CBA assays. The CBA assay, performed using a kit (BD Biosciences), allows the simultaneous detection and quantification of soluble murine IFN-γ, IL-17A, and IL-17F in a single sample. The principle of the CBA assay is as follows: five bead populations of equal size but distinct fluorescence intensities (resolved in the FL3 channel of a flow cytometer) are coated with capture antibodies specific for the different cytokines and mixed to form the CBA. The cytokine capture beads are mixed with recombinant cytokine standards or test samples and then mixed with the PE-conjugated detection antibodies, which are resolved in the FL2 channel. Following acquisition of sample data by flow cytometer, the results were analyzed using the BD CBA analysis software. Standard curves, plotted using the MFI of the beads for cytokine values ranging from 0 to 5,000 pg/ml, were used for quantifying the sample cytokines.

### Quantification of PtdInsP(3,4,5)_3_/PtdInsP(4,5)_2_

The levels of C18:0/C20:4-PtdIns(4,5)P_2_ and C18:0/C20:4-PtdIns(3,4,5)P_3_ in naïve CD4^+^ T cells were measured by using a liquid chromatography-tandem mass spectrometry (Nexera system, LCMS-8050, Shimadzu, Kyoto, Japan). We followed the extraction procedure and derivatization method described in Clark *et al* (Clark, Anderson et al. 2011). 1 µL of the resulting cell extract (200 µL) was injected onto a C4 column (UPLC BEH C4, 1.7 μm particle size, 300 Å pore size, 100 mm length x 1 mm inner diameter, Waters, Ireland). Solvent A consisted of deionized water containing 0.1% (v/v) formic acid. Solvent B consisted of 0.1% (v/v) formic acid in acetonitrile. We optimized the gradient conditions for the chromatography: 0 – 3.0 min 60% B, 3.0 – 5.0 min linear gradient 100% B, 5.0 – 7.0 min 100% B, and re-equilibration at 60% B for 2.5 min (100 µL/min flow rate). C17:0/C20:4-PtdIns(3,4,5)P_3_ standards (Avanti Polar Lipids, Alabaster, Alabama, USA) were used as an internal standard. Synthetic C18:0/C20:4-PtdIns(4,5)P_2_ and C18:0/C20:4-PtdIns(3,4,5)P_3_ standards (Avanti Polar Lipids) were used to determine the retention time (RT), the *m*/*z* values of the precursor ions, and the MS/MS fragmentation patterns.

### Mitochondrial respiration and glycolysis assay

CD4^+^ T cells (2 × 10^5^) were seeded in a 96-well plate and were stimulated with 2.5 μg/ml of anti-CD3 antibody (clone: 145-2C11) and 0.5 μg/ml anti-CD28 antibody (clone: 37.51). After two days of culture, cell pellets were collected for assay. Mitochondrial respiration and glycolysis were determined by measuring OCRs and ECARs in the extracellular space with a Seahorse XF96 Extracellular Flux Analyzer with the XF Cell Mito Stress Test Kit and XF Glycolysis Stress Test Kit (Seahorse Bioscience) according to the manufacturer’s instructions.

### Quantification and statistical analysis

Data analysis was performed using GraphPad Prism Software 7.0 (La Jolla, CA). Statistics were calculated using unpaired Student’s *t*-test or Mann-Whitney test. Means are given as ± SEM, with *P* values considered significant as follows: # < 0.05, \**P* < 0.05, \*\**P* < 0.01, \*\*\**P* < 0.001, and \*\*\*\**P* < 0.0001.

## Author Contributions

C.M.Y., D.K., S.H., R.H.S., S.K., and S.H.L. conceived the project and designed the experiments. C.M.Y., D.K., S.H., S.K., and S.H.L. executed most of the experiments, interpreted the data, and helped with the discussion of the results. M.K., S.J.P., H.M., and W.K. helped with *in vivo* experiments and gave advice. H.W.J. provided data analysis and helped with the discussion of the results. C.M.Y., D.K., S.H., M.K., S.K., and S.H.L. wrote the manuscript. R.H.S., S.K., and S.H.L. supervised all experiments and participated in the discussion of all results. S.H.L. had final responsibility for the decision to submit for publication.

## Ethics declarations

The authors declare no competing interests.

## Acknowledgements.

We thank Dr. Hyunji Lee for technical assistance, and the Hyewha Forum and Dr. Suk-Jo Kang for helpful discussion. This study was supported by grants funded by the Korea Mouse Phenotyping Project (NRF-2014M3A9D5A01073789 to R.H.S.), the National Research Foundation of Korea (NRF-2018R1A5A1024261 and NRF-2020R1A2C3005765 to S.K.), and the Biomlogic/GenoFocus (to S.H.L.).

