## Supplementary Table 1 for "Inositol polyphosphate multikinase regulates Th1 and Th17 cell differentiation by controlling Akt-mTOR signaling"

***Supplementary Table 1*—key resources table**

| **Antibodies** | **Source** | **Identifier** |
|---|---|---|
| phospho-STAT3 | 9131 | Cell Signaling Technology |
| STAT3 | 9139 | Cell Signaling Technology |
| phospho-STAT5 | 4322 | Cell Signaling Technology |
| STAT5 | 9363 | Cell Signaling Technology |
| T-bet | 14-5825-82 | eBioscience |
| GATA3 | sc-22206 | Santa cruz biotechnology |
| Rorγt | sc-293150 | Santa cruz biotechnology |
| GAPDH | Sc-32233 | Santa cruz biotechnology |
| β-actin | 4970 | Cell Signaling Technology |
| Anti-mouse phospho mTOR – percp-e710 | 46-9718-41 | eBioscience |
| Anti-mouse CD126 - PE | 12-1261-80 | eBioscience |
| Anti-mouse CD25 - PE | 12-1251-82 | eBioscience |
| Anti-mouse CD28 (clone 37.51) | 16-0281-86 | eBioscience |
| Anti-mouse CD3e (clone 145-2C11) | 16-0031-86 | eBioscience |
| Anti-mouse TCRβ – Percp-cy5.5 | 45-5961-82 | eBioscience |
| Anti-mouse CD4 – e450 | 48-0042-82 | eBioscience |
| Anti-mouse CD4 – PE/cy7 | 25-0041-82 | eBioscience |
| Anti-mouse CD44 – APC/cy7 | 47-0441-82 | eBioscience |
| Anti-mouse CD44 – APC | 17-0441-82 | eBioscience |
| Anti-mouse CD62L - FITC | 11-0621-82 | eBioscience |
| Anti-mouse CD62L – PE | 12-0621-82 | eBioscience |
| Anti-mouse CD8 – APC/cy7 | 47-0081-82 | eBioscience |
| Anti-mouse Foxp3 – APC | 17-5773-82 | eBioscience |
| Anti-mouse IFN-γ – FITC | 11-7311-82 | eBioscience |
| Anti-mouse IL-17A – APC | 17-7177-82 | eBioscience |
| Anti-mouse IL-17F – Percp-cy5.5 | 46-7471-82 | eBioscience |
| Anti-mouse IL-10 – FITC | 11-7101-82 | eBioscience |
| Anti-mouse T308 – PE | 558275 | BD biosciences |
| Anti-mouse S473 – APC | 17-9715-42 | eBioscience |
| Anti-mouse S6 – PE/cy7 | 25-9007-42 | eBioscience |
| Anti-mouse IFN-γ mAb functional grade | 16-7311-85 | eBioscience |

| | | |
|---|---|---|
| Anti-mouse IL-4 mAb functional grade | 16-7041-85 | eBioscience |
| Anti-mouse IL-12 mAb functional grade | 16-7123-85 | eBioscience |
| Anti-mouse IPMK | | Provided from Solomon Snyder Lab |
| **Chemicals, peptides, and recombinant proteins** | | |
| DPBS | LB001-02 | Welgene |
| percoll | 9139 | GE Heathcare |
| RPMI 1640 | LM011-01 | Welgene |
| RPMI 1640 nophenol | 11835030 | Gibco |
| IMDM | 12440061 | Gibco |
| BrefeldinA solution (1000X) | 00-4506-51 | eBioscience |
| PMA | 16561-29-8 | Calbiochem |
| Ionomycin | I24222 | ThermoFisher Scientific |
| Fixable Viabillity Dye eFlour 506 | 5536 | eBioscience |
| Ack lysis buffer | 2972 | Gibco |
| Foxp3/Transcription Factor staining buffer set | 2280 | eBioscience |
| IC Fixation buffer | 82-0000-49 | eBioscience |
| MOG peptide 35-55 | 2217 | Peptron |
| Complete Freund's adjuvant | F5881-6X | Sigma Aldrich |
| Pertussis toxin | 181 | List Biological |
| Recombinant mouse IL-2 | 212-12 | Peprotech |
| Recombinant mouse IL-12 | 210-12 | Peprotech |
| Recombinant mouse IL-4 | 404-ML | R&D SYSTEMS |
| Recombinant mouse IL-1β | 575106 | BioLegend |
| Recombinant mouse IL-23 | 589004 | BioLegend |
| Recombinant mouse IL-6 | 575706 | BioLegend |
| Recombinant mouse TGF-β | 100-21C | Peprotech |
| **Critical Commercial Assays** | **Source** | **Identifier** |
| Mouse naive CD4 T isolation Kit | 8804-6824-74 | eBioscience |
| CellTrace violet cell proliferation kit | C34557 | Invitrogen |
| CBA Mouse IL-17A flex set | 560283 | BD biosciences |
| CBA Mouse IFN-γ flex set | 558296 | BD biosciences |
| RNeasy mini kit | 74104 | QIAGEN |
| FastStart Universal SYBR Green Master (Rox) | 04913850001 | Roche |

| | | |
|---|---|---|
| **LC/MS** | | |
| ACQUITY UPLC Protein BEH C4 Column | 186005590 | Waters |
| ACQUITY ULPC Protein BEH C4 VanGuard Pre-Column | 186004623 | Waters |
| Trimethylsilyl diazomethane 2M solution in hexanes | 385330050 | Acros organics |
| C18:0/C20:4-PtdIns(3,4,5)$P_3$ | 850166 | Avanti Polar Lipids |
| C18:0/C20:4-PtdIns(4,5)$P_2$ | 850165 | Avanti Polar Lipids |
| C17:0/C20:4 PI(3,4,5)$P_3$ | LM1906 | Avanti Polar Lipids |
| **Experimental Models: Organisms/Strains** | | |
| C57BL/6J | | The Jackson Laboratory |
| B6.Cg-Tg(CD4-cre)1Cwi/BfluJ | JAX:022071 | The Jackson Laboratory |
| IPMK floxed mouse | This study | |
| **Oligonucleotides** | | |
| IPMK fwd: TGA AGA TTG GGC GGA AGA GC | This study | N/A |
| IPMK rev: GCC ATT GTG GAA AAA CTT GG | This study | N/A |
| Bactin fwd: GACAGGATGCAGAAGGAGATTAC | This study | N/A |
| Bactin rev: GCTGATCCACATCTGCTGGAA | This study | N/A |
| Hif1a fwd: CGGCGAAGCAAAGAGTCTG | This study | N/A |
| Hif1a rev: ATAACTGATGGTGAGCCTCATAAC | This study | N/A |
| Tbx21 fwd: GCCAGGGAACCGCTTATATG | This study | N/A |
| Tbx21 rev: GACGATCATCTGGGTCACATTGT | This study | N/A |
| Gata3 fwd: TTTACCCTCCGGCTTCATCCTCCT | This study | N/A |
| Gata3 rev: TGCACCTGATACTTGAGGCACTCT | This study | N/A |
| Rorc fwd: TGCAGGAGTAGGCCACATTAC | This study | N/A |
| Rorc rev: CCGCTGAGAGGGCTTCAC | This study | N/A |
| Rora fwd: CTCCCTGCGCTCTCCGCAC | This study | N/A |
| Rora rev: TCCACAGATCTTGCATGGA | This study | N/A |
| Foxp3 fwd: CCCAGGAAAGACAGCAACCTT | This study | N/A |
| Foxp3 rev: TTCTCACAACCAGGCCACTTG | This study | N/A |
| Il17a fwd: CTCCAGAAGGCCCTCAGACTAC | This study | N/A |
| Il17a rev: GGGTCTTCATTGCGGTGG | This study | N/A |
| Il23r fwd: GCCAAGAGAACCATTCCCGA | This study | N/A |
| Il23r rev: TCAGTGCTACAATCTTCAGAGGACA | This study | N/A |

| | | |
|---|---|---|
| Ccr6 fwd: AGGACTGGAGCCTGGATAACCAC | This study | N/A |
| Ccr6 rev: TAGGGCTTGAGATGATGATGGAGA | This study | N/A |
| IRF4 fwd: CAAGCAGGACTACAATCGTGAGGA | This study | N/A |
| IRF4 rev: AGTAGGAGGATCTGGCTTGTCGAT | This study | N/A |
| Il17f fwd: CCCATGGGATTACAACATCACTC | This study | N/A |
| Il17f rev: CACTGGGCCTCAGCGATC | This study | N/A |
| Ifng fwd: GGCCATCAGCAACAACATAAGCGT | This study | N/A |
| Ifng rev: TGGGTTGTTGACCTCAAACTTGGC | This study | N/A |
| Tnf fwd: CCACCACGCTCTTCTGTCTA | This study | N/A |
| Tnf rev: GATCTGAGTGTGAGGGTCTGG | This study | N/A |
| Csf2 fwd: CTAACATGTGTGCAGACCCG | This study | N/A |
| Csf2 rev: GTCTGGTAGTAGCTGGCTGTC | This study | N/A |
| Slc2a1 fwd: GAGACCAAAGCGTGGTGAGT | This study | N/A |
| Slc2a1 rev: GCAGTTCGGCTATAACACTGG | This study | N/A |
| Socs3 fwd: AGTGCAGAGTAGTGACTAAACATTACAAGA | This study | N/A |
| Socs3 rev: AGCAGGCGAGTGTAGAGTCAGAGT | This study | N/A |
| **Software and Algorithms** | | |
| Flowjo v.10.6.0 | FlowJo,LLC | http://www.flowjo.com |
| Prism 7 | GraphPad | http://www.graphpad.com |
| ImageJ | NIH | https://imagej.nih.gov/ij/ |
| Image Lab v.6.0.0 | Bio-Rad laboratories | https://www.bio-rad.com |
| MeV_4_8 | | http://mev.tm4.org/ |
