## Supplementary material for "Inositol polyphosphate multikinase regulates Th1 and Th17 cell differentiation by controlling Akt-mTOR signaling": Figure supplement

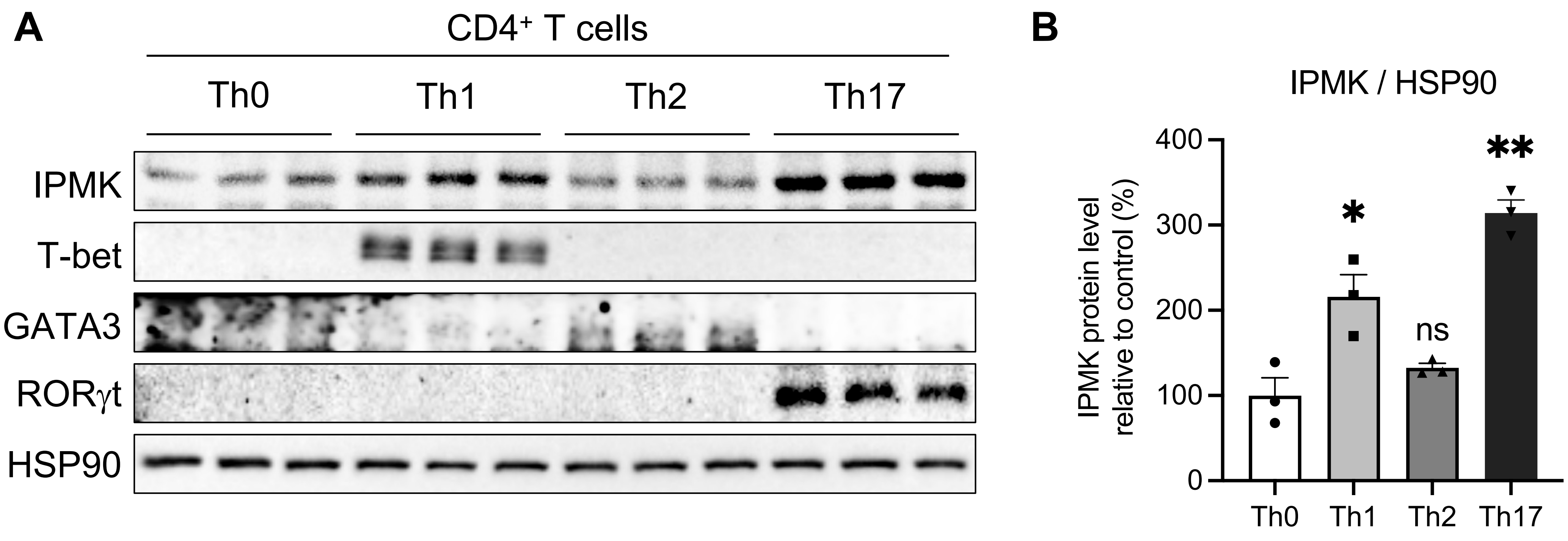


**Figure supplement 1.** Expression of IPMK in effector CD4 T cells. (**A and B**) Immunoblotting to detect IPMK protein expression in effector CD4 T cells activated under indicated differentiated conditions for 3 days. Results are representative of three independent experiments. Unpaired Student's t test was used for statistical analysis; *****$p$ < 0.05 and ******$p$ < 0.01 compared with the Th0 group.

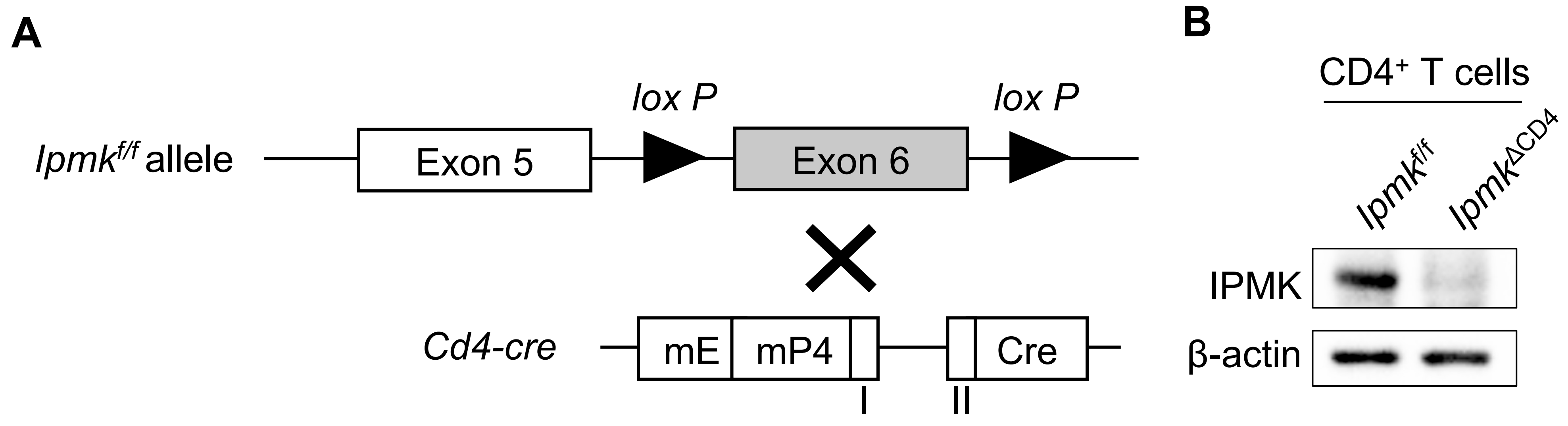


**Figure supplement 2.** Genetic deletion of IPMK in T cells. (**A**) Diagram depicting the structure of *Ipmk* gene showing exon 5 and 6, and the site of lox P insertion nearby exon 6 of *Ipmk*. Floxed IPMK mice without cre expression as controls (*Ipmk*$^{f/f}$) and crossed with cre recombinase driven by the *Cd4* promoter (it consists of the murine proximal enhancer (mE), The CD4 promoter (mP4) and a part of intron1). (**B**) Immunoblot analysis of IPMK expression in flow cytometry sorted CD4$^{+}$ T cells from IPMK floxed and Ipmk$^{f/f}$-CD4-Cre mice.

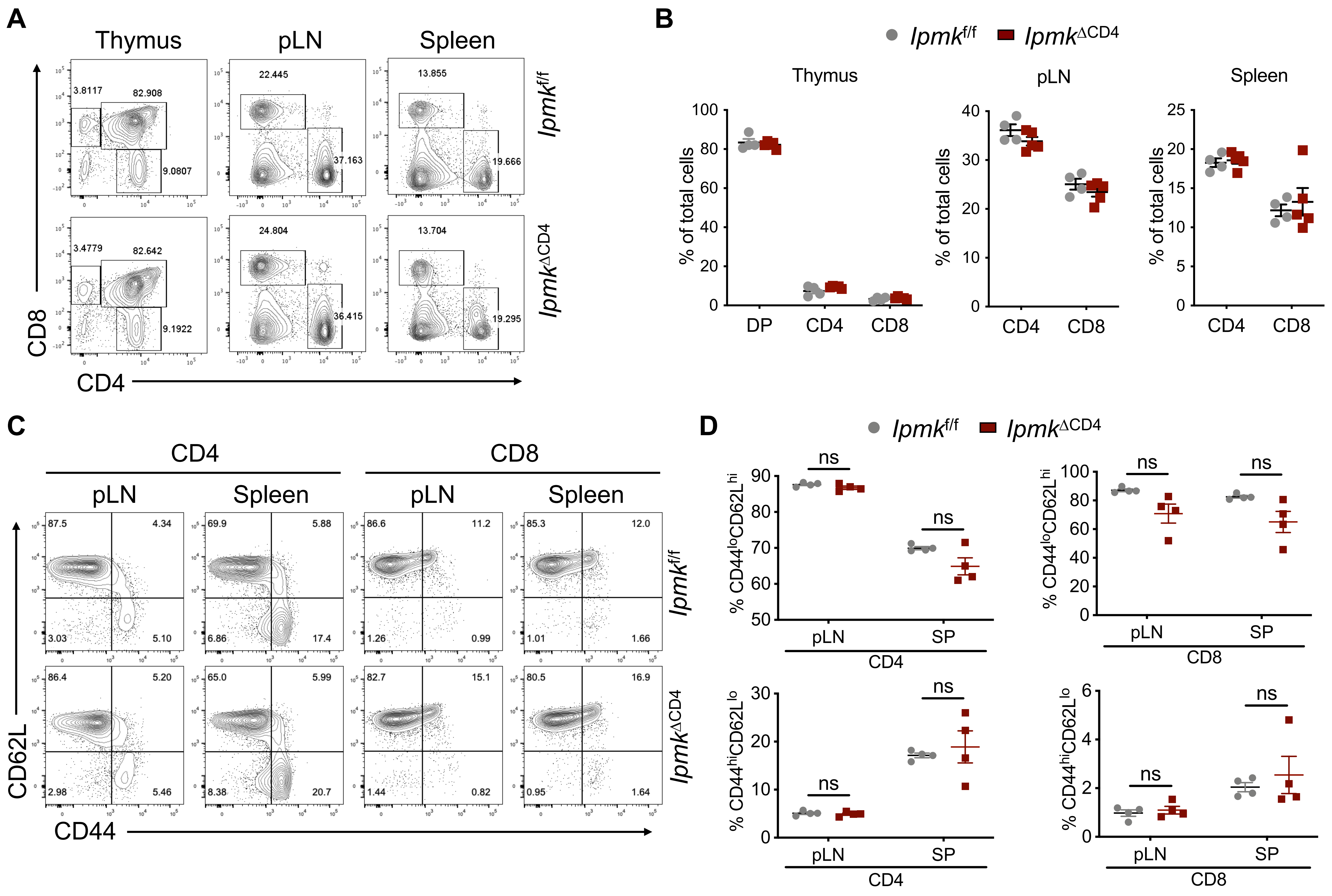


**Figure supplement 3.** The development of T cells in *Ipmk*$^{\Delta CD4}$ mice. (**A and B**) Cell surface staining profiles of CD4 or CD8 of *Ipmk*$^{f/f}$ or *Ipmk*$^{\Delta CD4}$ thymocytes, splenocytes or lymphocytes. The summary data for the cell surface staining profiles of CD4 or CD8 in thymocytes and splenocytes from *Ipmk*$^{f/f}$ or *Ipmk*$^{\Delta CD4}$ mice (n=4 or n=5 mice). (**C and D**) Cell surface staining of CD44 and CD62L expression on CD4 or CD8 T cells from pLN and spleens from *Ipmk*$^{f/f}$ or *Ipmk*$^{\Delta CD4}$ mice (n=4 mice; ns, not significant, per Mann Whitney test; centers indicate the mean values). Each figure shows representative and compiling data. Data are presented as the mean ± SEM; Unpaired Student's t test was used for statistical analysis; ns (not significant) compared with the *Ipmk*$^{f/f}$ group.

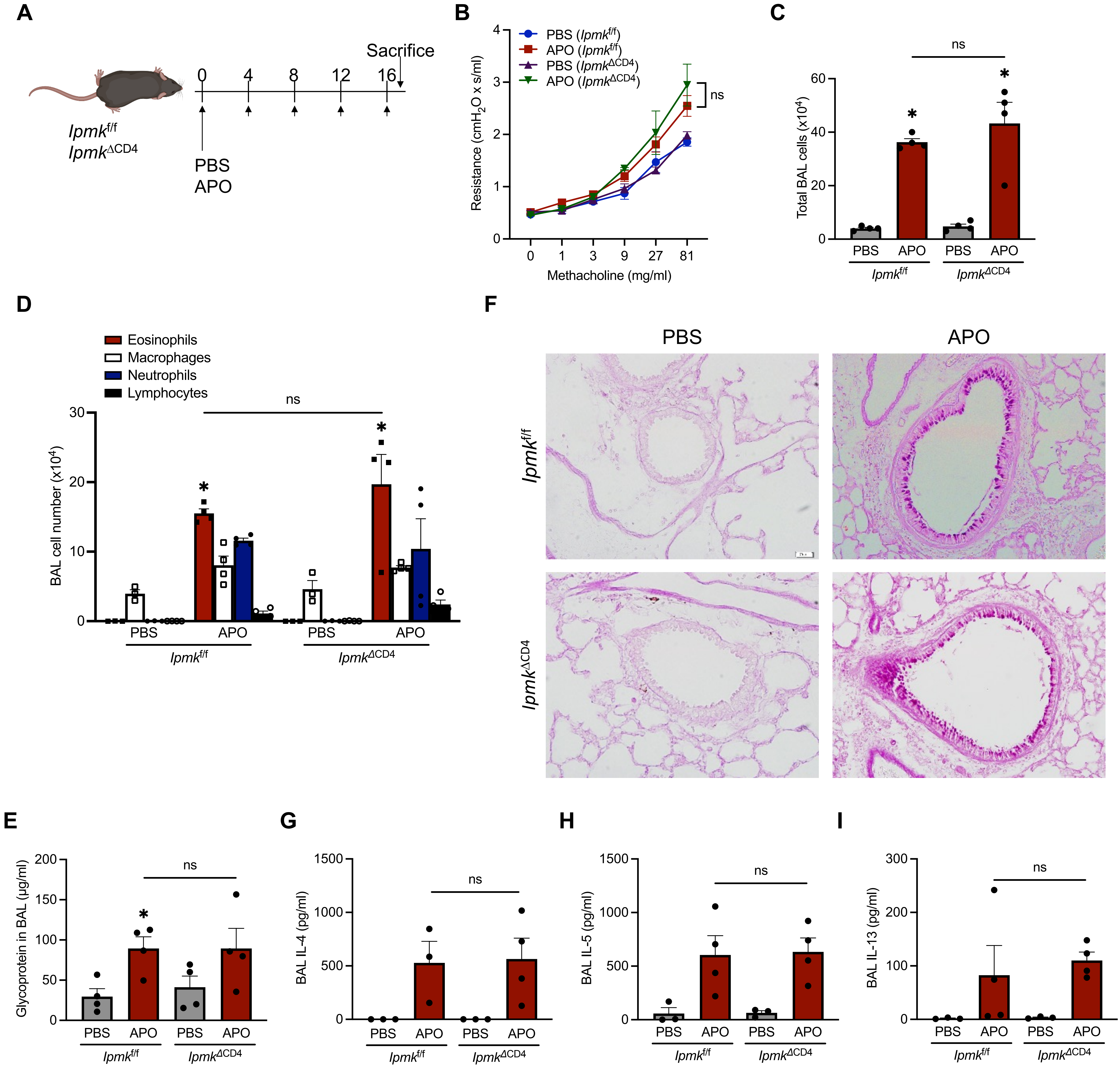


**Figure supplement 4.** Normal Th2-mediated immune responses without IPMK in CD4 T cells. (**A**) Schematic diagram for the induction of allergic asthma. (**B**) Airway hyperresponsiveness (AHR) was measured using the flexiVent system following by methacholine challenge [*Ipmk*$^{f/f}$ PBS (●), *Ipmk*$^{f/f}$ APO (■), *Ipmk*$^{\Delta CD4}$ PBS (▲), and *Ipmk*$^{\Delta CD4}$ APO (▼)]. (**C-E**) Total cell number (**C**), differential cell counts (**D**, ■: eosinophils, □: macrophages, ■: neutrophils, ■: lymphocytes), and glycoprotein secretion (**E**) in bronchoalveolar lavage fluid (BALF) were determined. (**F**) PAS staining of lung cross-sections was shown. (**G-I**) Production of Th2-associated cytokines BALF. Concentrations of IL-4 (**G**), IL-5 (**H**), and IL-13 (**I**) in BALF were measured by ELISA. Data are presented as the mean ± SEM; **p* < 0.05 and ns (not significant) compared with the each control group; by the Mann-Whitney test.

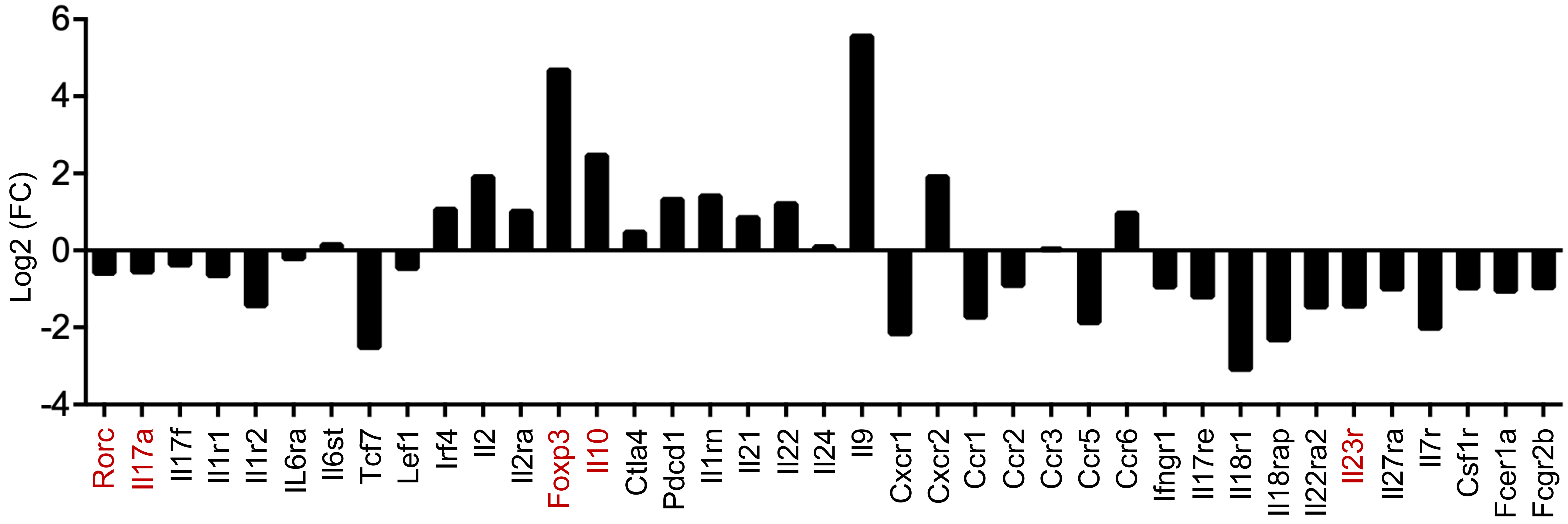


**Figure supplement 5.** List of genes differently expressed between *Ipmk*$^{f/f}$ or *Ipmk*$^{\Delta CD4}$ cells. The log2 fold change from *Ipmk*$^{f/f}$ mice of genes associated with the Th17 differentiation program are shown (n=2).
